# Characterization of mice with cell type-specific *Gnal* loss of function provides insights on *GNAL*-linked dystonia

**DOI:** 10.1101/2025.07.02.662743

**Authors:** Sophie Longueville, Ruiyi Yuan, Claire Naon, Emmanuel Valjent, Assunta Pelosi, Emmanuel Roze, Louise-Laure Mariani, Jean-Antoine Girault, Denis Herve

## Abstract

Isolated dystonia can be caused by loss-of-function mutations in the *GNAL* gene (DYT-GNAL). This gene encodes the α_olf_ heterotrimeric G protein subunit, which, together with β_2_γ_7_ subunits, mediates the stimulatory coupling of dopamine D1 and adenosine A2A receptors to adenylyl-cyclase. These receptors are expressed in distinct striatal projection neurons (SPNs) with complementary functions on motor behavior. To dissect the specific roles of Gα_olf_ in each subpopulation of SPNs, we generated and characterized mouse models in which *Gnal* was conditionally deleted in neurons expressing either D1 receptors (D1-SPNs) or A2A receptors (A2A-SPNs). Our results confirmed the critical role of Gα_olf_ in regulating adenylyl-cyclase 5 and its coupling with D1 and A2A receptors. Mice with a selective loss of Gα_olf_ in D1-SPNs showed nocturnal hyperactivity, deficits in motor performances, but no overt abnormal movements or generalized motor disability. Our experiments also revealed that Gα_olf_ in D1-SPNs is not systematically required for locomotor responses induced by D1 agonists or psychostimulants. Selective loss of Gα_olf_ in A2A-SPNs did not affect motor abilities nor learning. However, this loss strikingly increased spontaneous locomotor activity that was not further enhanced by psychostimulant drugs (cocaine, D-amphetamine, methylphenidate) or a selective A2 agonist, KW6002, and was paradoxically reduced by caffeine Our study identified specific roles of Gα_olf_ downstream of D1 and A2A receptors in the control of motor behavior and drug responses, highlighting their respective individual contribution in diseases associated with dysfunctional striatal signaling, including dystonia.

## 1. Introduction

Dystonia is a heterogeneous neurological entity characterized by sustained or intermittent abnormal movements and/or postures that are typically patterned and repetitive (Albanese et al., 2025). Dystonia has multiple causes and a broad phenomenological spectrum from adult-onset focal dystonia to infant-onset generalized dystonia, including tremulous and jerky forms. Converging evidence emphasizes the important role of dysfunctions of striatum in many patients with dystonia (Bhatia and Marsden, 1994; Neychev et al., 2011). More specifically, numerous defects of the dopamine-related striatal neurotransmission have been associated with dystonia (Ribot et al., 2019). They can be due to disorders disrupting dopamine synthesis or transport, altering the development or survival of dopaminergic neurons, or modifying the intracellular signal transduction downstream of dopamine receptors (Mencacci et al., 2020). Alterations in intracellular signaling neuron-specific in the striatum, particularly those involved in dopamine transmission, can cause various hyperkinetic motor disorders with dystonic components. These disorders may results from pathogenic variants in several genes, including *ADCY5* encoding adenylyl-cyclase 5 (AC5)(Chang et al., 2016; Chen et al., 2014), *PPP1R1B* encoding dopamine and cAMP-regulated phospho-protein 32 kDa (DARPP-32)(Khan et al., 2022), *GNAO1* encoding α_o_ subunit of G protein (Ananth et al., 2016) or several genes encoding phosphodiesterases (Erro et al., 2021; Miyatake et al., 2018; Niccolini et al., 2018; Salpietro et al., 2018). *GNAL*-linked dystonia (DYT-*GNAL),* typically a form of adult-onset isolated dystonia, is caused by pathogenic variants of *GNAL* (Fuchs et al., 2013; Goodchild et al., 2013), a gene encoding the Gα_olf_ subunit highly expressed in striatal neurons (Drinnan et al., 1991; Herve et al., 1993). The Gα_olf_ subunit has a high homology with the ubiquitously expressed Gα_s_ protein with the similar ability to stimulate adenylyl-cyclase (AC) (Jones and Reed, 1989; Liu et al., 2001; Xie et al., 2015). In the principal neurons of the striatum, the striatal projection neurons (SPNs), Gα_olf_ is much more expressed than Gα_s_ and thus plays a crucial role in the positive coupling of dopamine D1 (D1R) and adenosine A2A (A2AR) receptors with AC (Corvol et al., 2001; Zhuang et al., 2000). In some populations of GABAergic interneurons and in cholinergic interneurons Gα_olf_ and Gα_s_ are co-expressed (Herve et al., 2001; Millett et al., 2024). Careful analysis of the functions of Gα_olf_ proteins carrying pathogenic variants has shown the diversity of mechanisms leading to the disruption of G protein-coupled receptor (GPCR) signaling (Masuho et al., 2018). However, variants generating truncated and non-functional forms of Gα_olf_ can cause DYT-*GNAL* dystonia (Fuchs et al., 2013; Kumar et al., 2014). Therefore, mouse models in which a variant in *Gnal* sequence interrupts Gα_olf_ expression would correctly mimic the alterations present in some patient families. These models have a substantial limitation because homozygous invalidation of *Gnal* (*Gnal*-/-) leads to a very high lethality in the first days after birth, caused in part by anosmia, which disrupts feeding reflexes in newborns (Belluscio et al., 1998). Heterozygous *Gnal* KO rodents (*Gnal*+/-) are fully viable and reproduce the genetic condition seen in human disease, in whom most pathogenic variants act in a dominant manner (Masuho et al., 2018). While both mice and rats do not show spontaneous abnormal movements reminiscent of dystonia, they may present alterations in motor coordination and motor learning (Pelosi et al., 2017; Yu-Taeger et al., 2020). In *Gnal*+/- mice, abnormal movements are triggered by oxotremorine, a muscarinic agonist, revealing their vulnerability to dystonic-type manifestations (Pelosi et al., 2017). Although Gα_olf_ is expressed outside the striatum, particularly in motor regions such as the cerebellum and cortex (Millett et al., 2024; Vemula et al., 2013), its specific role in striatal dysfunction appears important in DYT-*GNAL*. A recent study has shown that the selective conditional *Gnal* KO in the striatum at embryonic or adult stages leads to abnormal postures, reminiscent of human symptoms. In the striatum, there are two main classes of SPNs. One class expresses the dopamine D1R and forms the direct pathway (dSPNs) projecting directly to the basal ganglia output structures, namely the *internal globus pallidus* and the *substantia nigra pars reticulata*. The other class expresses dopamine D2 receptors (D2Rs) and A2ARs, and initiates several relays in the basal ganglia before reaching the output structures (indirect pathway iSPNs) (Gerfen and Surmeier, 2011). The two classes of SPNs have complementary functions, with dSPNs stimulating spontaneous and acquired motor behaviors and iSPNs inhibiting competing motor actions, which promote the emergence of context-adapted responses (Durieux et al., 2009; Hikida et al., 2010; Kravitz et al., 2012; Nelson and Kreitzer, 2014). These processes may be disrupted in dystonia leading to maladaptive motor command and, subsequently, abnormal movements and postures (Mink, 2003). In this study, we address the role of Gα_olf_ in each specific subpopulation of SPN on mouse motor behavior to better understand the pathophysiology of DYT-*GNAL*. We also explore how its absence in dSPNs or iSPNs affects responses to drugs acting on dopamine or adenosine signaling to test their possible effects in this disease.

## 2. Materials and Methods

### 2.1 Generation of *Gnal* conditional KO mice

A targeting vector with the exons 3 and 4 of *Gnal* flanked by LoxP sites was introduced by homologous recombination in C57BL/6N embryonic stem cells and chimeric mice were produced at Phenomin (Strasbourg, France). The male chimeric mice were bred with female *Flp* deleter mice to excise the FRT-flanked selection cassette (FRT-Neo-FRT) and generate F1 heterozygotes with the floxed *Gnal* allele (*Gnal*^f/+^). To generate conditional KO mice used in this study, *Gnal*^f/f^ homozygotes were bred to either Drd1Cre (Tg(Drd1-cre)EY262^Gsat^) (Gong et al., 2007) or Adora2aCre (Tg(Adora2a-cre)^2MDkde^) (Durieux et al., 2009) mice, expressing Cre recombinase under the control of D1R (*Drd1*) and A2AR (*Adora2a*) gene promoters, kindly provided by Pr Paul Greengard (Rockefeller University) and Dr Alban de Kerchove d’Exaerde (Université libre de Bruxelles), respectively. These crosses generated *Gna*l-D1cKO and *Gnal*-A2AcKO mice with *Gnal* knockout in neurons expressing D1R and A2AR, respectively. Mice bearing *Gnal*^f/+^ genotype and Cre-expressing transgene were crossed with *Gnal*^f/f^ mice to obtain male and female experimental animals. For genotyping, DNA was extracted from a tail biopsy taken between postnatal days 4 and 7, and processed for PCR using primers to test the Cre transgene (Nestin CreER F: ATTTGCCTGCATTACCGGTC; Nestin CreER R: ATCAACGTTTTCTTTTCGG, annealing temperature 57°C) as well as floxed and wild-type *Gnal* (Gnal-Lf : CCACAGTGTCAAACTGCCTGTATCGC Gnal-Lr : CAAGGCATCTGCCAAGTTTTTACCC; annealing temperature 62°C). PCR fragments were resolved in 2 or 3% agarose gel. In several litters, pup survival and weight were assessed daily. In all experiments, mice with the *Gnal*^f/+^ or *Gnal*^f/f^ genotypes that lack the Cre transgene were considered as wild-type mice (WT) because Gα_olf_ expression in *Gnal*^f/f^ mice was not different from that in *Gnal*^+/+^ mice (see below).

The mice were maintained in a 12-h light/dark cycle, in stable conditions of temperature (21 ± 1°C), with access to food and water *ad libitum* and housed 2 to 6 per cage. All animal procedures used in the present study were approved by the Ethical committee for animal experiments (CEEA Charles Darwin) and by the *Ministère de l’Education Nationale de l’Enseignement Supérieur de la Recherche, France* (projects APAFIS #8196 and #37798). The laboratory animal facility was approved to carry out animal experiments by the *Sous-Direction de la Protection Sanitaire et de l’Environnement de la Préfecture de Police* (licence D75-05-22).

### 2.2 Drugs

Selective A2AR antagonist KW6002 (istradefylline, #5147), non-selective adenosine antagonist caffeine (#2793), selective A2AR agonist CGS21680 hydrochloride (#1063), and the selective D1R agonists, SKF81297 hydrobromide (#1447), dihydrexidine hydrochloride (#0884) and, SKF83822 hydrobromide (#2075) were purchased from Tocris. Cocaine hydrochloride was provided by Cooper (Melun, France) and D-amphetamine sulfate and methylphenidate hydrochloride by Sigma. KW6002, SKF81297, dihydrexidine, SKF83822 and CGS21680, used for investigating their effects on locomotion activity, were dissolved in a solution containing 5% dimethylsulfoxide (DMSO), 5 % Tween20, 15 % polyethylene glycol (both vol/vol) in H_2_O. Caffeine, D-amphetamine, cocaine and methylphenidate were dissolved in 9 g L^-1^ NaCl. For cAMP assays, SKF81297 hydrobromide (0.1 mM) was dissolved in H_2_O and CGS21680 hydrochloride (30 mM) in DMSO. Drugs were then diluted in H_2_O to obtain the appropriate concentrations.

### 2.3 Western blot analysis

Mice were killed by decapitation and their brain was quickly dissected out, frozen in 2-methylbutane (Sigma) cooled at −30°C in dry ice and then stored at −80°. Olfactory bulbs were removed and frozen in dry ice. Using a Cryostat (Leica), frozen brains were cut into 210 µm-thick slices from which the dorsal striatum, nucleus accumbens and frontal cortex were punched out using cold stainless-steel cannulas (1.4 mm for dorsal striatum and nucleus accumbens, and 2.2 mm for frontal cortex). The samples were sonicated in 10 g L^-1^ sodium dodecyl sulfate (SDS) and heated at 100°C for 5 min. Aliquots (5 μl) of homogenate were used for measuring protein concentration using a bicinchoninic acid (BCA) assay kit (Thermo technologies). Equal amounts of protein (20 μg) were separated by 15–10% (w/v) pre-casted gel (Bio-Rad) in the presence of SDS and transferred to nitrocellulose membranes (Trans-Blot Turbo Transfer, Bio-Rad). The membranes were immunoblotted using the following primary antibodies: rabbit polyclonal affinity-purified Gα_olf_ and Gα_s_ (1:1000)(Herve et al. 2001), AC5 (Millipore MAPS2049, 1:500), mouse monoclonal DARPP-32 (1:1000) (gift of Pr P. Greengard), Gβ_2_ (Sigma AV48278, 1:1000), rabbit polyclonal Gγ_7_ (1:1000)(gift of Pr Robishaw), mouse monoclonal D1R (1:1000) (gift of Pr Luedtke), chicken polyclonal TH (Aves, #TYH, 1:1000), D2R (Frontier Institute, D2-Rb-Af960, 1:500), A2AR (Frontier Institute, A2A-Go-Af700, 1:500) and mouse or rabbit polyclonal actin (Sigma-Aldrich, A5441, A2066, 1:5000) and IRDye 800CW-or 700CW-conjugated anti-mouse, anti-chicken, anti-goat and anti-rabbit IgG (1:5000) (Rockland Immunochemical) as secondary antibodies. Their binding was quantified using an Odyssey–LI-COR infrared fluorescent detection system (LI-COR).

### 2.4 Measures of AC activity

Mice were killed by cervical dislocation and the brains were quickly removed from the skull. The right and left striata were dissected with glass manipulators on ice and homogenized in 600 µL of a cold buffer containing 2 mM Tris-maleate (pH 7.2), 2 mM EGTA and 300 mM sucrose, using a Potter-Elvehjem apparatus. AC activity was measured by incubating tissue homogenates (6-12 µg of protein) for 10 min at 30°C in 30 µL of a reaction mix containing 25 mM Tris-maleate (pH 7.2), 0.5 mM ATP, 1 mM MgSO4, 0.33 mM EGTA, 0.1 mM papaverine, 2 µg.mL^-1^ adenosine deaminase, 10 mM creatine phosphate, 0.3 mg mL^-1^ creatine kinase, 1 µM GTP and 50 mM sucrose. For every homogenate, AC activities were determined in the presence of vehicle or various concentrations of SKF81297 or CGS21680. The reaction was stopped by adding 20 µL of cold lysis buffer (Cisbio kit) and cooling the tube on ice. The levels of cAMP formed were determined in 1/20 of reaction volume by cAMP – Gs Dynamic kit according to the manufacturer’s protocol (Cisbio Bioassays). Förster resonance energy transfer (FRET) in plate wells was quantified (330 nm excitation, 620 nm and 665 nm emission of respective fluorophores) and analyzed using a Homogeneous Time Resolved Fluorescence (HTRF) program on a Misthras LB940 instrument (Berthold). The ratio obtained from Signal 665 nm/Signal 620 nm was used to interpolate cAMP concentrations from a cAMP standard curve obtained from the same plate and calculated with GraphPad Prism (GraphPad Software, La Jolla, CA). Protein concentration in tissue homogenates was measured using a bicinchoninic acid (BCA) assay (Micro BCA protein assay kit, Thermo Scientific). AC activity was expressed in pmoles of cAMP formed per min per mg of protein and normalized as % of basal activity in WT type striatum.

### 2.5 Behavioral tests

Behavioral experiments were conducted using mutant and control littermates of both sexes, aged 2-8 months, between 8 am and 6 pm.

#### Locomotor activity

Locomotor activity was measured in a circular corridor (or circular maze) with four infrared beams placed at every 90° angle (Imetronic, Pessac, France) in a low luminosity environment. Counts were incremented by consecutive interruptions of two adjacent beams (1/4 turn). Mice were habituated to handling 3 days before testing. In one experiment, animals were placed in the circular maze at 11 am, for 28 hours to assess their circadian rhythm of activity with free access to water and food throughout the test. In other experiments to test drug responses, mice were habituated to the device during 3 sessions of 90 min (1 session/day). On day 1, they were just placed in the maze. On days 2 and 3, they additionally received an intraperitoneal (ip) injection of saline 30 min after the start of the recording. We used the locomotor activity during the first 30 min (before saline injection) to evaluate response to a novel environment and habituation. The following days, drug response was assessed by comparing locomotor activities after vehicle and drug administrations on two consecutive days. Recording lasted 90 or 120 min depending on the drug tested, and vehicle and drug were administered ip 30 min after the start of the recording. The drug doses used were selected based on previously published studies: 5 mg.Kg^-1^ SKF81297 and dihydrexidine (Yano et al., 2018), 0.5 mg.Kg^-1^ SKF83822 (Brunori et al., 2021; O’Sullivan et al., 2004); 0.5 mg.Kg^-1^ CGS21680 (Ledent et al., 1997), 3.3 mg.Kg^-1^ KW6002 (Xiao et al., 2006), 15 mg.Kg^-1^ caffeine (El Yacoubi et al., 2000), 20 mg.Kg^-1^ cocaine (Corvol et al., 2007), 2.5 mg.Kg^-1^ D-amphetamine and 10 mg.Kg^-1^ methylphenidate (Pascoli et al., 2005).

#### Rotarod

Mutant and WT littermates of either sex were trained for 7 days on the rotarod (BIOSEB, model LE8200), with 7 trials per day, spaced by 5-min intervals using a procedure adapted from (Varani et al., 2024). The trial started by placing mice on a rod with a rotation accelerating from 4 to 40 rpm over 5 min and ended when either the mice fell off the rod or 300 s elapsed. During the 5-min intervals, animals were placed in home cages. Before and 45 min after the 7 trials, animals were placed in open field boxes (50 x 50 x 30 cm) made of white Plexiglas and allowed to freely explore it during 5 min. Locomotion was tracked using a ceiling-mounted camera (Panasonic WV BP332) and ViewPoint videotrack. The distance traveled was measured to verify the absence of gross locomotor deficit in the tested mice. The rotarod was cleaned using 30% ethanol (vol/vol) between subjects.

#### Pole test

Motor function was assessed using Pole test in mutant and WT littermates of either sex. The device consisted of a 50 cm high, 1 cm diameter vertical pole covered with elastic adhesive tape to allow the animals to grip it. The pole was placed in the middle of a cage. The mouse was placed at the top of the pole, head up, and the mouse turning and descent were videotaped. On the video films, we measured the time to turn completely downward (Tturn) and that to descend to the floor (Tdown). The maximum time for Tturn was set to 120 s. After this time, the mice were gently pushed by hand to begin the descent. The test consisted of 5 trials per animal and the mean values of Tturn and Tdown were measured. Since there was no significant difference of Tdown between groups of mutant and WT mice, we have only used results on Tturn. The pole was carefully cleaned after testing between mice.

#### Grid test

Animals were allowed to hang from a cage grid, then the grid was inverted and maintained about 30 cm above the home cage. The mice were videotaped and we measured the time (in s) it took the mice to fall off the grid. The average time in 3 trials was used to assess neuromuscular strength.

#### Grip test

Neuromuscular functions were also evaluated in groups of mutant and WT mice of either sex by determining the maximal peak force they developed when they were pulled out of a specially designed grid (Grip Strength Test, Bioseb). The grip strength measuring device was positioned horizontally. The animal, held by the tail, was able to grasp a metal grid with its front paws, and was then pulled backward in the horizontal plane. The force applied to the grid just before the mouse released its grip was recorded as maximal peak force by a force sensor. The results, expressed in grams (g), corresponded to the average of 5 consecutive trials. To account for size differences between the mice, the average maximum force (in g) was divided by body weight (in g).

#### Thigmotaxis analysis in open-field

White plastic squared boxes (50 x 50 x 40 cm) were used in the open-field test. Mice were initially placed in the center of the apparatus, allowed to freely explore the environment for 20 min. Their behavior was recorded by a camera positioned overhead. A central area was defined by a 25 × 25 cm square in the middle of the box. Time, distance, and velocity in the center and periphery were automatically analyzed with the Viewpoint software. The percentage of the distance in the central area relative to the total distance was used to evaluate avoidance of the center (thigmotaxis) of animals.

#### Elevated O-maze

The device consisted of a circular ring platform (O-maze, width 5 cm, outer diameter 51 cm) made of gray plastic, placed 40 cm above the ground, with 2 open quadrants without borders and 2 quadrants closed by an opaque sidewall of 15 cm, opposite to each other. Mice started the test on a closed quadrant, the animals facing an open quadrant, opposite to the experimenter. The test lasted 5 min and was recorded by a camera positioned above the experimental set up. The time spent in open and close quadrants and total traveled distance were automatically analyzed with the Viewpoint software. The time spent in the open quadrants was used to measure risk assessment in mice.

### 2.6 Statistical analysis

All statistical analyses were performed using GraphPad Prism 5 (GraphPad Software). Data are expressed as mean ± SEM. p values < 0.05 were considered to be statistically significant. All studies involved balanced groups of male and female mice and data from both sex were analyzed together. For responses of AC activity to various concentrations of D1R or A2AR agonists, the dose response curves were fitted using GraphPad Prism software (log(agonist) vs response). The responses in WT and mutant homogenates were compared with 2-way ANOVA. For results of Western blot analysis and some behavioral tests, we used the parametric statistic test, one-way ANOVA followed by Bonferroni’s post hoc test when normal distribution of the data was confirmed by the Shapiro–Wilk test. If not, non-parametric tests, Mann-Whitney test or Kruskal-Wallis test followed by Dunn’s multiple comparison test were applied. Data containing two variables (e.g., genotype and treatment, or treatment and time) were analyzed by two-way ANOVA followed by Bonferroni’s multiple comparison test. Repeated-measures two-way ANOVA was generally used when the mouse behavior was followed over time or the responses to drug and vehicle was compared in the same mice. We generated Kaplan-Meier plots with GraphPad Prism 5 for comparing the survival of pups with various genotypes obtained from crosses between *Gnal*-D1cKO+/- and *Gnal*^f/f^ mice and the difference significance was evaluated by the Log-rank (Mantel-Cox) test.

## 3. Results

### 3.1 *Gnal* KO in D1-SPNs and A2A-SPNs respectively impairs cAMP production in response to D1 and A2 agonists

We first checked that the introduction of LoxP sites into introns 2 and 4 in *Gnal*^f/f^ mice did not modify the amount of Gα_olf_ in the striatum (Gα_olf_/actin in % of the WT mean ± SEM; *Gnal*^f/f^, 90.5 ± 8.8; *Gnal*^+/+^, 100 ± 7.4, Mann-Whitney test, Mann-Whitney U = 13.0, p = 0.48). In the rest of the study, we used mice without Cre-expressing transgene (Drd1Cre or Adora2aCre) as controls, whether the *Gnal* gene was floxed or not, and they are designated as wild-type (WT). We compared AC activity in basal condition or in the presence of D1 or A2A agonist in striatal homogenates of *Gnal*^f/f^ x Drd1Cre/+ (referred to as *Gnal*-D1cKO-/-below) and WT mice. Basal AC activity was markedly reduced (by about 50%) in *Gnal*-D1cKO-/-mice. The D1 agonist, SKF81297, had no significant effect on AC activity while it strongly increased it in WT mice (**Fig. 1A**). These data showed that *Gnal* KO in D1-SPNs abolished the ability of D1Rs to activate AC. In contrast, the A2A agonist, CGS21680, stimulated AC activity in both *Gnal*-D1cKO-/- and WT striatum (**Fig. 1B**). CGS21680-stimulated activity in *Gnal*-D1cKO mice was lower than in WT mice but this appeared to essentially result from the decreased basal activity in these mice. When the response to CGS21680 was expressed as the difference between CGS21680-stimulated activity and basal activity, the maximal response in the presence of 10 µM CGS21680 was not different in *Gnal*-D1cKO-/- and WT mice (**Fig. 1C**).

**Fig. 1.**
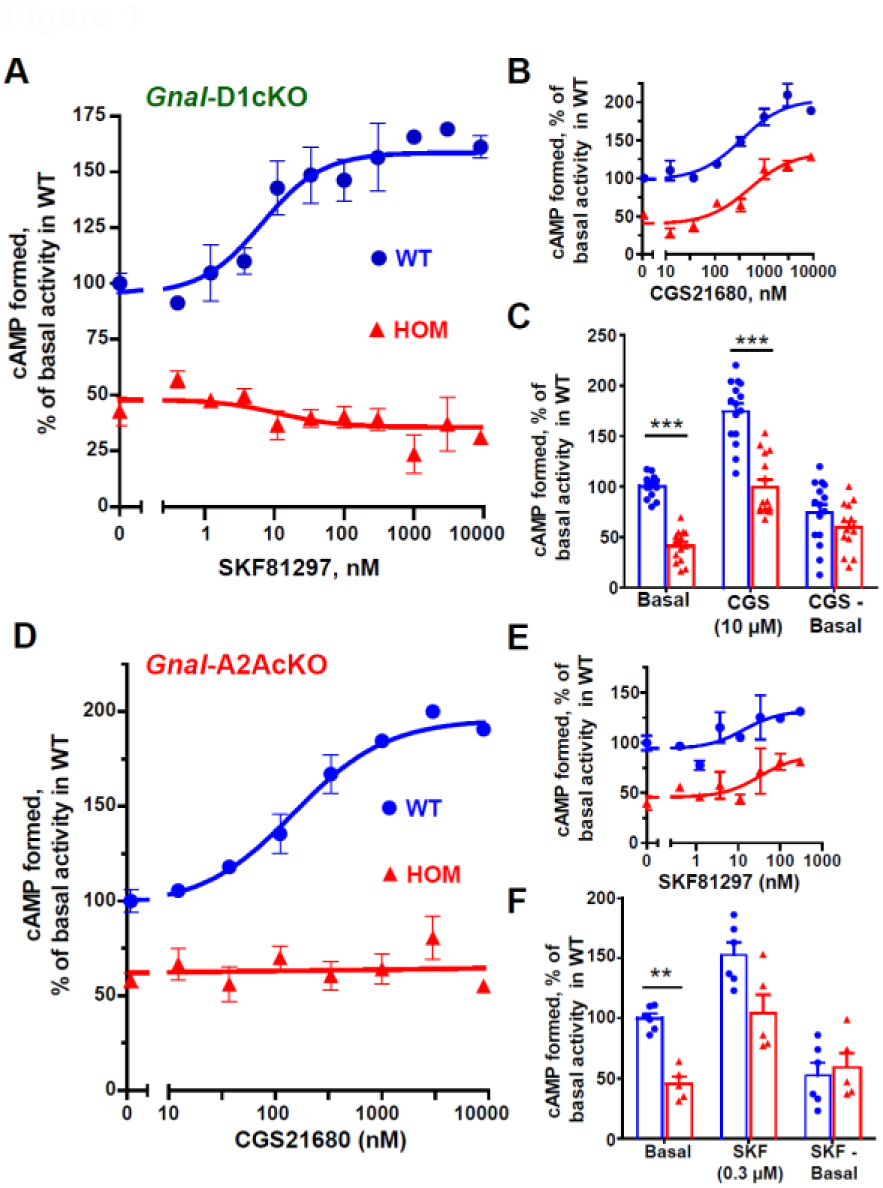
Adenylyl-cyclase responses to D1 or A2A agonists in the dorsal striatum of *Gnal* cKO mice. **A-B)** cAMP formation was measured in the presence or absence of various concentrations of the D1 agonist SKF81297 (A) or A2A agonist CGS21680 (**B**) in striatal homogenates from Gnal-D1cKO-/- (HOM) mice and wild-type (WT) littermates. In Gnal-D1cKO-/- mice, the adenylyl-cyclase response to D1 agonist was abolished (**A**), while that to A2A agonist remained significant (**B**). **C**) cAMP production under basal conditions or in the presence of a maximum concentration of CGS 21680 (10 μM) was reduced in mutant mice compared to WT littermates but the difference between CGS-stimulated and basal productions (CGS - basal) was not significantly different. **D-E**) cAMP formation was measured in the presence or absence of various concentrations of A2A agonist CGS21680 (**D**) or D1 agonist SKF81297 (**E**) in the striatal homogenates of Gnal-A2AcKO-/- (HOM) and WT mice. In Gnal-A2AcKO-/- mice, the adenylyl-cyclase response to A2A agonist was abolished, while that to D1 agonist remained significant. **F**) cAMP production under basal conditions or in the presence of a maximum concentration of SKF81297 (0.3 μM) was reduced in mutant mice compared to WT littermates but the difference between CGS-stimulated and basal productions (CGS - basal) was not significantly different. The data of cAMP production were expressed as % of basal activity evaluated in WT mice and are pooled from 3 (A, B) or 2 (E, F) separate experiments. Basal adenylyl-cyclase activity was 106 ± 32 (A-C) or 86 ± 18 (D-F) pmoles of cAMP formed.min-1.mg-1 Prot. All the statistical results are shown in Supplementary Statistical tables. The dose-response curves were best-fitted with GraphPad Prism software. Responses obtained in mutant and WT mice were compared with two-way ANOVA. ** p < 0.01 and ***, p < 0.001, significantly different from WT littermates.

In striatal homogenates of *Gnal*^f/f^ x Adora2aCre/+ mice (referred to as *Gnal*-A2AcKO-/-below), a significant reduction in basal AC activity and a complete absence of response to CGS21680 were observed, whereas AC activity was strongly increased by increasing concentrations of CGS21680 in WT homogenates (**Fig. 1D**). These data showed that homozygous *Gnal*-A2AcKO completely abolished the positive coupling of A2AR to AC. In contrast, in *Gnal*-A2AcKO-/-mice, the AC response to SKF81297 was preserved (**Fig. 1E**). The maximal response expressed as the difference between the activity in the presence of high concentration of SKF81297 (0.3 µM) and the basal activity did not show any difference between *Gnal*-A2AcKO and WT mice (**Fig. 1F**). All these data demonstrated that homozygous *Gnal* D1cKO and A2AcKO exclusively blocked the coupling of D1R and A2AR to AC, respectively.

### 3.2 Impact on proteins involved in intracellular signaling of conditioned *Gnal* silencing

In *Gnal*-D1cKO-/-mice, Gα_olf_ levels were strongly reduced in the dorsal striatum (**Fig. 2**) and *nucleus accumbens* (**Fig. S1A**). In *Gnal*-D1cKO+/-mice (Gnal^f/+^ x Drd1Cre/+), Gα_olf_ levels were also significantly decreased in both structures, but less strongly than in homozygotes (**Figs. 2** and **S1A**). Because of its role in olfaction, we also measured Gα_olf_ levels in olfactory bulbs where it is highly enriched (Herve et al., 1993; Yu-Taeger et al., 2020). We did not observe any decrease (**Fig. S1B**), ruling out potential alterations in olfaction in these mice. In SPNs, the Gα_olf_ subunit is predominantly in interaction with the G protein subunits Gγ_7_ and Gβ_2_ as well as with the effector protein AC5 (Schwindinger et al., 2010; Xie et al., 2015). Gβ_2_ and Gγ_7_ levels were not significantly changed (**Fig. 2**). In contrast, AC5 levels were decreased in both *Gnal*-D1cKO-/- and *Gnal*-D1cKO+/- mice, the effect being more pronounced and significant in homozygotes (**Fig. 2**). Interestingly, there was a significant correlation between striatal Gα_olf_ and AC5 levels **Fig. S1C**). It suggests that the absence of Gα_olf_ destabilizes AC5 in D1-SPNs and leads to a decrease of its levels. This effect can account for the decrease in basal AC activity in *Gnal*-D1cKO mice (see **Fig 1A**). Levels of Gα_s_, a protein highly homologous to Gα_olf_, were increased in the dorsal striatum of *Gnal*-D1cKO-/- mice (**Fig. 2**). However, since Gα_s_ expression was at least 10 times lower than that of Gα_olf_, this increase only marginally compensated for the absence of Gα_olf_. D1R levels tended to increase in both *Gnal*-D1cKO-/- and *Gnal*-D1cKO+/- mice, but not significantly (**Fig. 2**). D2R and A2AR levels, did not change (**Fig. S2**). The fact that DARPP-32 and tyrosine hydroxylase levels were not modified (**Figs. 2** and **S1A**) strongly suggested that the number and trophicity of striatal SPNs and their dopamine innervation were not altered in mutant mice.

**Fig. 2.**
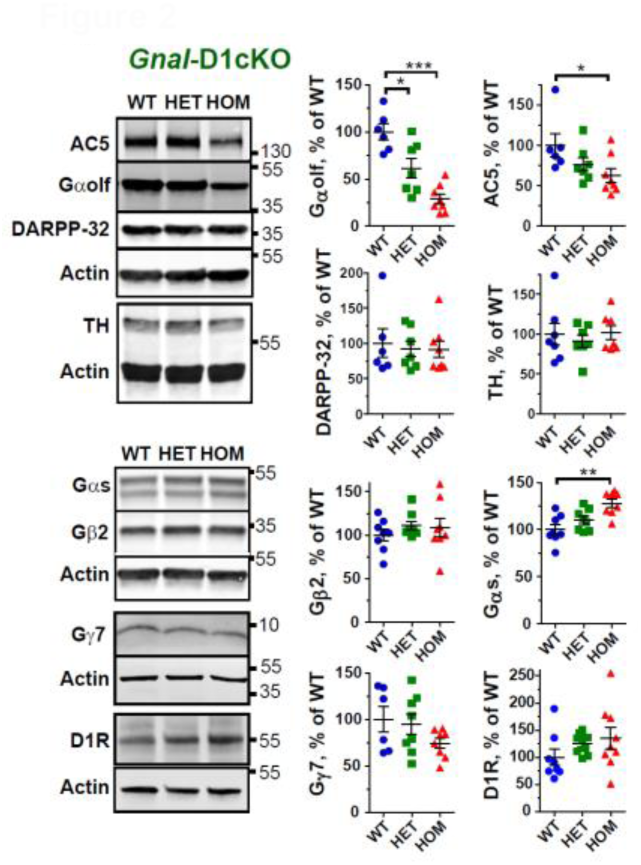
Molecular characterization of *Gnal*-D1cKO-/- and *Gnal*-D1cKO+/- mice. Representative immunoblots and quantification of Gαolf, adenylyl-cyclase 5 (AC5), DARPP-32, tyrosine hydroxylase (TH), β2 (Gβ2), γ7, (Gγ7) and αs (Gαs) subunits of G proteins and, D1 dopamine receptor (D1R) in dorsal striatum homogenates of *Gnal*-D1cKO-/- (HOM), *Gnal*-D1cKO+/- (HET) and wild-type (WT) littermates. Data corresponded to the ratio between the immunofluorescence of the indicated protein and actin, expressed as a % of the mean value in WT samples. Partial or complete loss of *Gnal* expression in D1-SPNs reduced significantly the levels of Gαolf, but also those of AC5. By contrast, no significant change was observed in Gβ2 and Gγ7 levels, two proteins associated with Gαolf. Some compensatory increase of Gαs could be detected. DARPP-32, D1R and TH (markers of SPNs, D1-SPNs and dopamine terminals, respectively) levels were not significantly altered. Protein levels were compared using one-way ANOVA followed by Bonferroni’s multiple comparison test or Kruskal-Wallis test followed Dunn’s multiple comparison test depending on whether data distribution was normal or not (see the results in Supplementary Statistical tables). * p<0.05, ** p < 0.01 and ***, p < 0.001.

In *Gnal*-A2AcKO-/- and *Gnal*-A2AcKO+/- (*Gnal*^f/+^ x Adora2aCre/+) mice, Gα_olf_ levels were decreased, the effect being more marked in homozygotes than in heterozygotes (**Fig. 3**). We also observed decreases in AC5 levels that can account for the reduction of baseline AC activity measured in striatal tissue homogenates (see **Fig. 1D**). D2R and A2AR levels tended to slightly decrease (**Fig. S3**). No changes in D1R, Gα_s_ and Gβ_2_ levels were observed (**Fig. 3**). The lack of alteration in DARPP-32 and tyrosine hydroxylase levels indicated that SPNs and striatal dopamine innervation were not grossly altered in *Gnal*-A2AcKO mice.

**Fig. 3.**
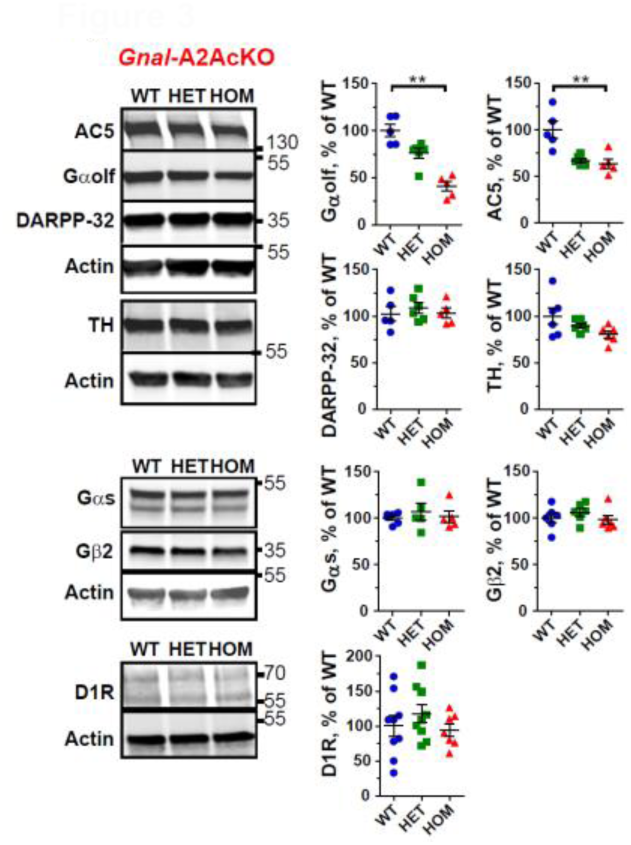
Molecular characterization of *Gnal*-A2AcKO-/- and *Gnal*-A2AcKO+/- mice. Representative immunoblots and quantification of Gαolf, adenylyl-cyclase 5 (AC5), DARPP-32, tyrosine hydroxylase (TH), β2 (Gβ2) and αs (Gαs) subunits of G proteins and, D1 receptor (D1R) in dorsal striatum homogenates of *Gnal*-A2AcKO-/- (HOM), *Gnal*-A2AcKO+/- (HET) and wild-type (WT) littermates. Data corresponded to the ratio between the immunofluorescence of the indicated protein and actin (Act), expressed as a % of the mean value in WT samples. Partial or complete loss of Gnal expression in D1-SPNs reduced the levels of Gαolf, but also those of AC5. By contrast, no significant change was observed in levels of G protein subunits, Gβ2 and Gαs levels. DARPP-32, D1R or TH (markers of SPNs, D1-SPNs and dopamine terminals, respectively) levels were not significantly altered. Protein levels were compared using one-way ANOVA followed by Bonferroni’s multiple comparison test or Kruskal-Wallis test followed Dunn’s multiple comparison test depending on whether data distribution was normal or not (see the results in Supplementary Statistical tables). ** p < 0.01, significantly different.

### 3.3 Selective *Gnal* KO in D1-SPNs impairs survival of pups at weaning

Genotyping of pups from crosses between *Gnal*-D1cKO-/- and *Gnal*^f/f^ mice, a few days after birth (4-7 days), excluded significant in utero or early postnatal mortality of mice lacking Gα_olf_ in D1-SPNs, as there was no significant bias from the expected Mendelian distribution (90 *Gnal*^f/+^, 91 *Gnal*^f/f^, 98 *Gnal*-D1cKO+/-, 70 *Gnal*-D1cKO-/-, χ2 = 0.173, NS). In contrast, we observed an increase in lethality in *Gnal*-D1cKO-/- from 21 days, around the weaning period, until the 40^th^ day (**Fig. 4A**). Ultimately, 34% of *Gnal*-D1cKO-/- pups died before age 60 days. When we retrospectively examined the body weight curve of pups that died before 60 days (**Fig. 4B**), we observed, at around 15 days, a halt in weight gain and even a drop in some individuals, suggesting a feeding defect. Interestingly, the other part of the population of *Gnal*-D1cKO-/- animals, surviving beyond 60 days, showed normal weight gain. When we determined the weight of adult mice (median age: females, 100 days; males, 122 days), we observed no significant difference between the different genotypes in either females or males (**Fig. 4C**). These data indicated that *Gnal*-D1cKO-/- animals surviving beyond 60 days had a normal development and maintained a body weight similar to that of WT mice, suggesting a normal feeding.

**Fig. 4.**
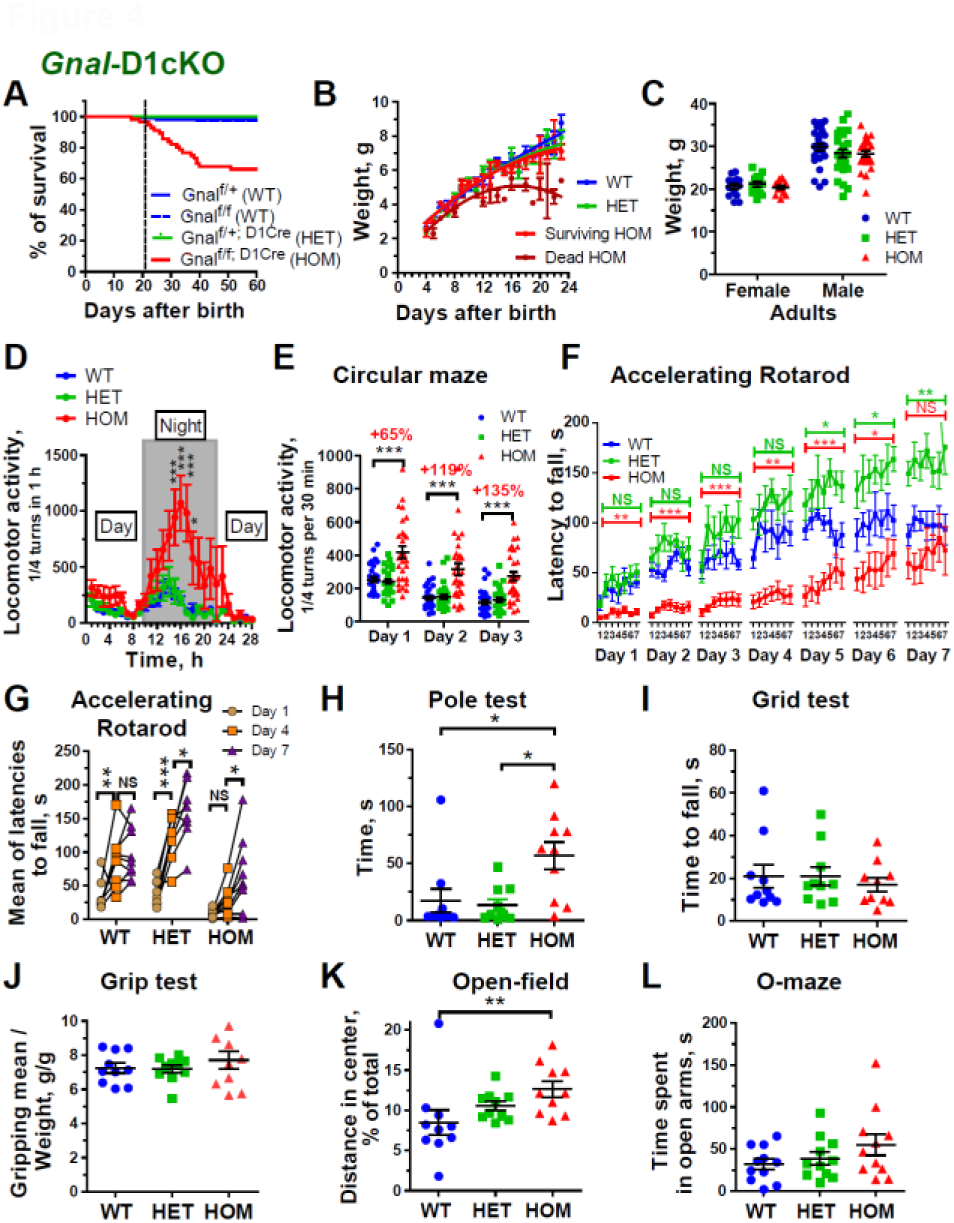
Survival and motor behavior of *Gnal*-D1cKO-/- and *Gnal*-D1cKO+/- mice. **A)** Survival curve of mice from the same litters with different genotypes within the first 60 days after birth. A portion of *Gnal*-D1cKO-/- (HOM) mice died around and after postnatal day 21, corresponding to the weaning period (vertical dashed line indicates day 21). This was not the case in littermates with other genotypes (wild-type, WT and *Gnal*-D1cKO+/-, HET). The difference was significant according to log-rank (Mantel-Cox) test (see **Supplementary Statistical tables**). **B)** Weight curve of mice from the same litters with different genotypes. In *Gnal*-D1cKO-/- mice, data for mice alive and dead at postnatal day 60 were separated. **C)** Weights of male and female adult mice (median age: 100 and 122 for females and males, respectively) with different genotypes. Data were analyzed by two-way ANOVA (Genotype and Sex factors, see **Supplementary Statistical tables**). **D)** Spontaneous locomotor activity on 28 h indicating a hyperactivity in *Gnal*-D1cKO-/- mice, the circadian rhythm remaining unaltered. **E)** Hyperactivity was also observed in these mutant mice when exploratory activity was assessed in a novel environment (circular maze) or during sessions on the 2 following days. In **D** and **E**, data were analyzed by repeated measures two-way ANOVA followed by Bonferroni’s multiple comparison test. *** p<0.001 (see **Supplementary Statistical tables**). **F)** Motor coordination and motor learning as measured by the Rotarod. Mean ± SEM of latency to fall (in seconds, s) in each session. A deficit in motor coordination was observed in *Gnal*-D1cKO-/- mice. Both homozygous and heterozygous mutant mice showed a progressive improvement in Rotarod performance over multiple sessions/days to the point that the performance of *Gnal*-D1cKO+/- mice became superior to that of WT mice on day 7. For each day, data were analyzed by repeated measures two-way ANOVA (Genotype and Trial factors) (see **Supplementary Statistical tables**). * p<0.05, ** p<0.01, *** p<0.001, genotype factor between mutant and WT littermates. NS, not significant. **G)** Motor learning on days 4 and 7 in the Rotarod. Each point is the average fall latency over 7 sessions. Data were analyzed by repeated measures two-way ANOVA (Genotype and Day factors) followed by Bonferroni’s multiple comparison test (see **Supplementary Statistical tables**). * p<0.05, ** p<0.01, *** p<0.001, different between days. **H** Motor coordination as measured by the pole test. Time to turn completely downward at the top of the pole (Tturn) was measured and each point is the mean of 3 trials. *Gnal*-D1cKO-/- mice showed a deficiency compared to WT littermates. **I** and **J**) Assessment of physical strength as measured by grid test and grip strength. **I**, the graph represents the time (in sec) it takes for mice to fall when they are caught on a grid. **J**, Each point corresponds to the average grip strength (maximal force in g, 5 trials/mouse) divided by the mouse weight (in g) to take in account difference in mouse size. There was no genotype difference in performances in these two tests. **K** and **L**, Evaluation of anxiety in mutant mice compared to WT littermates. In **K**, the % distance traveled in the center of an open field was measured to assess the ability to overcome anxiety. In **L**, the same parameter was evaluated by the time spent by mice in open arms of an elevated O-maze. *Gnal*-D1cKO-/- mice appeared less anxious than their WT littermates. **H-L**, data were compared using Kruskal-Wallis test followed Dunn’s multiple comparison tests (see **Supplementary Statistical tables**). * p<0.05, ** p<0.01.

### 3.4 Selective *Gnal* KO in D1-SPNs increases nocturnal locomotor activity and impairs motor performance in adults

*Gnal*-D1cKO-/- mice exhibited locomotor hyperactivity that was particularly marked during the night period (**Fig. 4D**). When they were tested during the day, *Gnal*-D1cKO-/- mice also showed increased activity when placed in the circular maze for 30 min (**Fig. 4E**). Although locomotor activity was reduced by daily repetition of the tests, it remained significantly increased compared to WT mice. *Gnal*-D1cKO+/- mice exhibited locomotor activities similar to those of WT mice.

We then tested motor performance and motor learning in *Gnal*-D1cKO-/-, *Gnal*-D1cKO+/- and WT mice in an accelerating Rotarod. The mice were tested for 7 days with 7 sessions per day (**Fig. 4F**). *Gnal*-D1cKO-/- mice had much lower performances than WT or *Gnal*-D1cKO+/- mice. However, as indicated by their locomotor hyperactivity, this deficiency could not be attributed to a generalized impaired ability to move. The disturbance in Rotarod performances of *Gnal*-D1cKO-/- mice remained significant until the 6^th^ day of testing, but we observed a progressive increase in performances. On day 7, their scores, although lower, did not differ significantly from those of WT mice (**Fig. 4F**). These *Gnal*-D1cKO-/- mice were therefore capable of motor learning. *Gnal*-D1cKO+/- mice, which had motor coordination performances similar to WT mice on day 1, showed remarkable learning abilities in the accelerating Rotarod. Their scores increased day by day to the point of exceeding those obtained by WT. Observation of *Gnal*-D1cKO+/- mice in the Rotarod showed them to be highly focused on the task, compared to WT. In this experiment, the scores of WT mice increased significantly between days 1 and 4, then plateaued thereafter (**Fig. 4G**). In contrast, in heterozygous and homozygous mice, performances increased significantly between days 4 and 7 (**Fig. 4G**).

Motor abilities of *Gnal*-D1cKO-/- mice were also evaluated using the Pole test. They took significantly longer time to turn around and get off the pole (**Fig. 4H**). Visual observations indicated that mutant mice had great difficulty initiating the relatively complex actions of turning around at the top of the pole. On the other hand, the descent was rapid. *Gnal*-D1cKO+/- mice performed like WT in this test (**Fig. 4H**). Interestingly, *Gnal*-D1cKO-/- and *Gnal*-D1cKO+/- mice showed no hindlimb clasping when suspended by the tail (**Fig. S4A**). In addition, visual observation of these mice did not indicate any obvious abnormalities in posture or spontaneous movements compared to WT mice. In tests that evaluate neuromuscular strength in mice such as the Grid and Grip tests (**Figs. 4I** and **4J**), there was no significant deficit in *Gnal*-D1cKO homozygotes, showing the absence of significant neuromuscular weakness that could explain the poor motor performances in the Rotarod and Pole tests.

In the Open field, *Gnal*-D1cKO-/- mice visited the central area of the arena more frequently than WT or *Gnal*-D1cKO+/- mice (**Fig. 4K**). However, the total distance traveled was similar in the three groups of mice (total distance as % of WT mean: *Gnal*-D1cKO-/-, 110 ± 12; *Gnal*-D1cKO+/-, 99 ± 7; WT, 100 ± 7; one-way ANOVA, F_(2, 27)_ = 0.43, NS). In this experimental condition in which locomotor activity was measured during 20 min in an open arena, we did not detect in the *Gnal*-D1cKO-/- mice the locomotor hyperactivity previously observed in a circular maze with a low-light environment (see above). The more frequent presence of *Gnal*-D1cKO-/- mice in the central area of the open field could indicate that these mice were less anxious than WT mice. However, when we subjected the mice to a test specifically evaluating risk assessment (elevated O maze, **Fig. 4L**), no significant difference could be detected between the different groups of mice. It indicated that the deficits found in the Rotarod in *Gnal*-D1cKO-/- mice were not related to elevated risk assessment.

### 3.5 Selective *Gnal* KO in A2A-SPNs results in high locomotor hyperactivity without impaired motor coordination

When we monitored the survival curve of litters from mating of *Gnal*-A2AcKO+/- mice with *Gnal*^f/f^ mice, we did not observe any particular mortality in the resulting genotypes (**Fig. 5A**). Adult *Gnal*-A2AcKO-/-male mice displayed a slight body weight deficit compared to the other groups of mice, but this deficit was not observed in females (**Fig. 5B**).

**Fig. 5.**
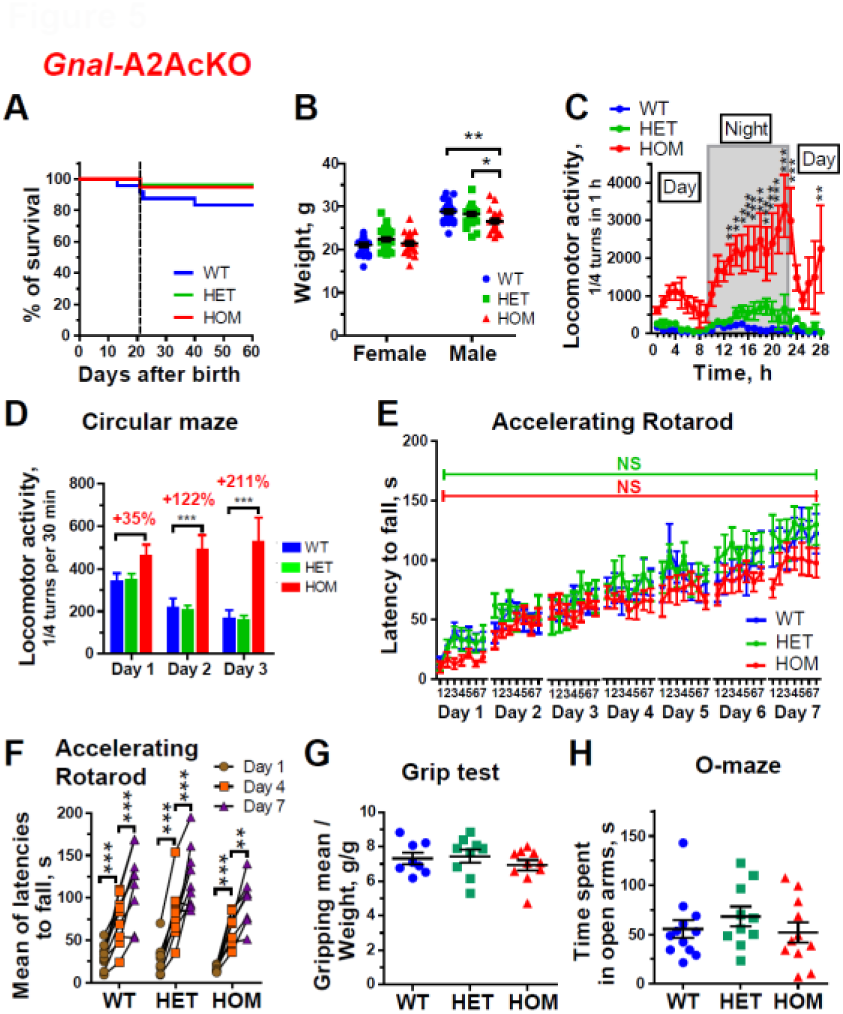
Survival and motor behavior of *Gnal*-A2AcKO-/- and *Gnal*-A2AcKO+/- mice. **A)** Survival curve of *Gnal*-A2AcKO-/- (HOM) and *Gnal*-A2AcKO+/- (HET) mice compared to wild-type (WT, *Gnal*f/f or *Gnal*f/+) from the same litters within the first 60 days after birth. Postnatal day 21 (vertical dashed line), corresponding to the weaning period. No significant difference was detected using log-rank (Mantel-Cox) test (see **Supplementary Statistical tables**). **B)** Weights of male and female adult mice (median age: 113 and 103 for females and males, respectively) with different genotypes. Data were analyzed by two-way ANOVA (Genotype and Sex factors) followed by Bonferroni’s multiple comparison test (see **Supplementary Statistical tables**). * p<0.05, ** p<0.01. **C)** Spontaneous locomotor activity on 28 h indicating a hyperactivity in *Gnal*-A2AcKO-/- mice, the circadian rhythm remaining unaltered. **D)** Bar graphs (mean ± SEM) indicating hyperactivity in these mutant mice when exploratory activity was assessed in a novel environment or during sessions on the 2 following days. In **C** and **D**, data were analyzed by repeated measures two-way ANOVA followed by Bonferroni’s multiple comparison test. ** p<0.01, *** p<0.001 (see **Supplementary Statistical tables**). **E)** Motor coordination and motor learning as measured by the Rotarod. Mean ± SEM of latency to fall (in s) in each session. No deficit in motor coordination was observed in HOM or HET mice compared to WT mice. NS, not significant. Both HOM and HET mutant mice showed a progressive improvement in Rotarod performance over multiple sessions/days similar to that of WT animals. For each day, data were analyzed by repeated measures two-way ANOVA (Genotype and Trial factors) (see **Supplementary Statistical tables**). NS, not significant. **F)** Motor learning on days 4 and 7 in the Rotarod. Each point is the average fall latency over 7 sessions. Data were analyzed by repeated measures two-way ANOVA (Genotype and Day factors) followed by Bonferroni’s multiple comparison test (see **Supplementary Statistical tables**). ** p<0.01, *** p<0.001, different between days. **G)** Assessment of strength as measured by grip test. Each point corresponds to the average grip strength (maximal force in g, 5 trials/mouse) divided by the mouse weight (in g) to take in account difference in mouse size. There was no significant difference between genotypes. **H)** To assess anxiety in mutant and WT mice, the ability to overcome anxiety was evaluated by the time spent by mice in open arms of an elevated O-maze. **G** and **H**, data were compared using Kruskal-Wallis test (see **Supplementary Statistical tables**).

When we examined locomotor activity in a circular corridor for 28 h, we noticed that *Gnal*-A2AcKO-/-mice exhibited a strong locomotor hyperactivity that was particularly apparent during the night phase (**Fig. 5C**). It is possible that the lower body weight of *Gnal*-A2AcKO-/-males resulted from an abnormally high physical expenditure in these mice. However, female *Gnal*-A2AcKO-/-mice appeared more hyperactive than males (**Fig. S5**) but this did not appear to have had any impact on their body weight, which was normal, similar to that of WT mice. When mice were placed every day during three consecutive days in a circular maze at daytime, WT and *Gnal*-A2AcKO+/-animals displayed an exploratory activity within the first 30 min of exposure in this novel environment on day 1, but gradually habituated to the circular maze so that their activity decreased on day 2 and 3 (**Fig. 5D**). In contrast, the activity of *Gnal*-A2AcKO-/-mice that was not significantly different from WT or *Gnal*-A2AcKO+/-mice on day 1 (**Fig. 5D**), increased progressively on day 2 and 3, when it was about 3-fold higher than in WT mice (**Fig. 5D**). This indicates that *Gnal*-A2AcKO-/-mice displayed a constant locomotor hyperactivity with a lack of habituation.

We then examined motor abilities and motor learning in the accelerating Rotarod using multiple sessions as described above (see 3.4). Both *Gnal*-A2AcKO-/- and *Gnal*-A2AcKO+/-mice increased their performance over time, showing motor learning similar to WT mice (**Fig. 5E**). In this experiment, the progression of performance was similar in the three genotypes (**Figs. 5E** and **5F**). Both *Gnal*-A2AcKO-/- and *Gnal*-A2AcKO+/-mice showed no overt abnormal movements or hindlimb clasping when suspended by the tail (**Fig. S4B**). Grip strength of mutant mice was not significantly different from that measured in WT mice (**Fig. 5G**). In the elevated O-maze, the time spent in the open arm was the same in heterozygous and homozygous mutant mice as in WT, suggesting that the hyperactivity of *Gnal*-A2AcKO-/-mice did not result from an altered risk assessment.

### 3.6 Selective *Gnal* KO in D1-SPNs reduces responses to D1 receptor agonists but does not alter responses to pharmacological modulators of A2A receptors

Because the absence of *Gnal* expression in D1-SPNs abolished AC response to D1R agonist in striatal homogenates, we investigated behavioral responses to two D1R agonists, SKF81297 and dihydrexidine. When locomotor activity was measured in response to SKF81297 (5 mg.Kg^-1^), responses of *Gnal*-D1cKO-/-mice were variable but reduced compared with those observed in WT mice (**Fig. 6A**). The locomotor response to dihydrexidine (5 mg.Kg^-1^) was completely lost in mice lacking Gα_olf_ in D1-SPNs (**Fig. 6B**). In *Gnal*-D1cKO+/-mice, the responses to dihydrexidine were normal. Surprisingly, locomotor response to another D1R agonist, SKF83822 (0.5 mg.Kg^-1^), which is reported to be specifically associated with AC activation (Undie et al., 1994), appeared not to depend on Gα_olf_ expression (**Fig. S6**).

**Fig. 6.**
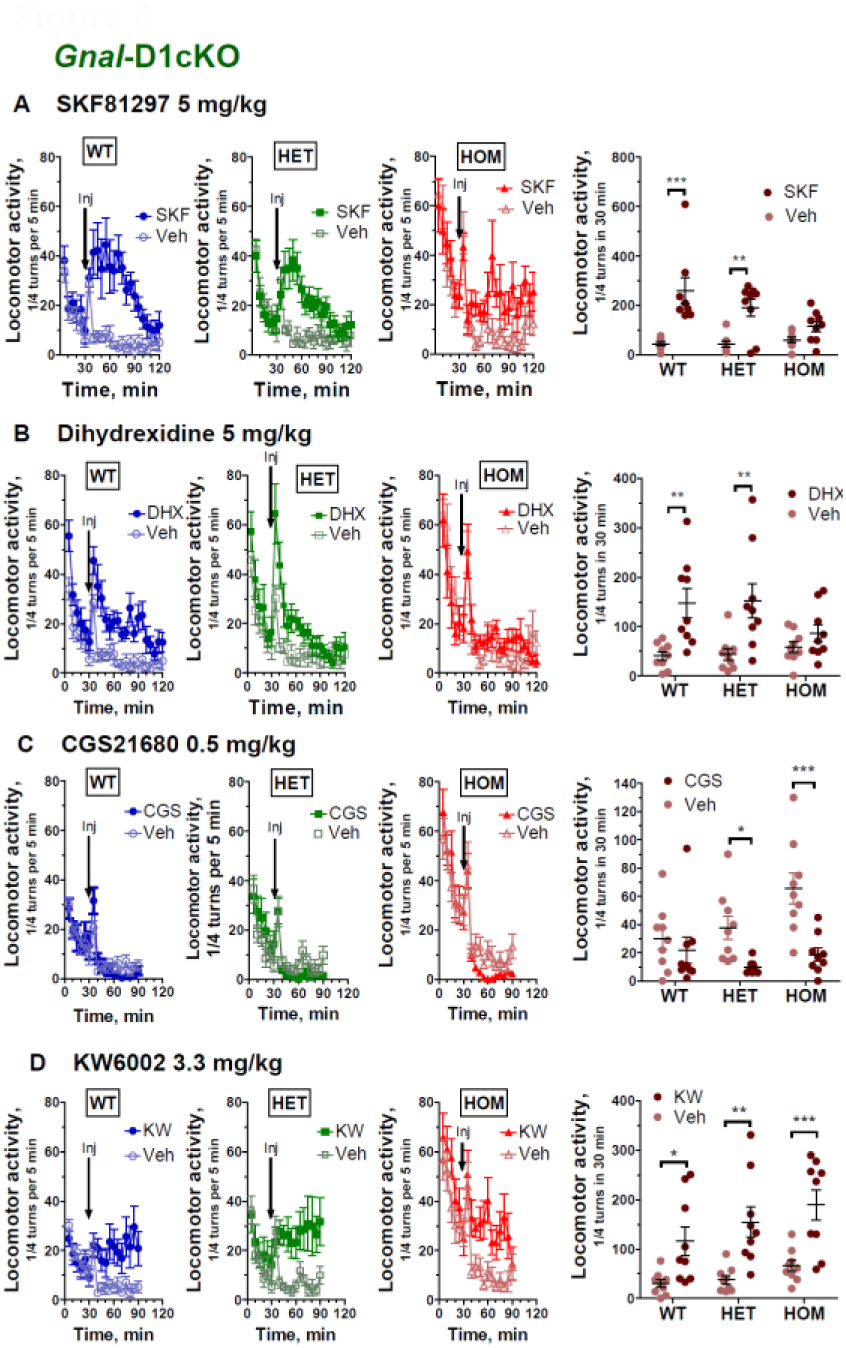
Effects of D1R or A2AR ligands on locomotion in *Gnal*-D1cKO-/- and *Gnal*-D1cKO+/-mice. **A-D)** time course of locomotor activity (3 left panels), before and after an ip injection of the D1R agonist SKF81297 (SKF, 5 mg.Kg-1) (**A**), D1R agonist dihydrexidine (DHX, 5 mg.Kg-1) (**B**), A2AR agonist CGS21680 (CGS, 0.5 mg.Kg-1) (**C**) or A2AR antagonist KW6002 (KW or istradefylline, 3.3 mg.Kg-1) (**D**) in wild-type (WT), *Gnal*-D1cKO+/-(HET) and *Gnal*-D1cKO-/-(HOM) mice. In all cases, comparisons were made with locomotion time course after vehicle administration. Each point is the mean ± SEM of the number of ¼ turns in a circular corridor during 5 min. Time-courses of responses to drug and vehicle were compared using repeated measures two-way ANOVA (Treatment and Time factors) followed by Bonferroni’s multiple comparison test (see **Supplementary Statistical tables**). **A-D**, right panels, cumulative responses on 30 min after drug or vehicle administration in WT, HET and HOM mice. Graphs represent scatter dot plots with mean ± SEM. Responses to drug and vehicle in mouse groups with different genotypes were compared using repeated measures two-way ANOVA (Genotype and Treatment factors) followed by Bonferroni’s multiple comparison test (see **Supplementary Statistical tables**). * p<0.05, ** p<0.01, *** p<0.001.

We also investigated whether locomotor responses to A2AR agents were altered in mice with reduced or no *Gnal* expression in D1-SPNs. The decrease in locomotor activity in response to administration of a selective A2AR agonist, CGS21680 (0.5 mg.Kg^-1^), which was very small (−27%) in WT mice, was significant in *Gnal*-D1cKO+/-(−74%) and *Gnal*-D1cKO-/-(−71%) mice (**Fig. 6C**). Response to A2AR agonist thus appeared to be somehow enhanced in these mice. When we treated mice with a selective A2AR antagonist, KW6002 (a.k.a. istradefylline, 3.3 mg.Kg^-1^), the increase in locomotor activity within 30 min of injection was similar in WT (+288%), *Gnal*-D1cKO+/-(+310%) and *Gnal*-D1cKO-/-(+288%) mice (**Fig. 6D**). These results clearly indicated that the absence of *Gnal* expression in D1-SPNs significantly reduced locomotor responses to D1R agonists but not to A2AR agonist or antagonist.

### 3.7 Selective *Gnal* KO in D1-SPNs impairs responses to psychostimulants

Psychostimulants increase dopamine, noradrenaline and serotonin in the synaptic extracellular space through various mechanisms (Docherty and Alsufyani, 2021). We tested the consequences of *Gnal* KO in D1-SPNs on responses to three psychostimulants, cocaine, D-amphetamine and methylphenidate. In *Gnal*-D1cKO+/-mice, the locomotor response to cocaine (20 mg.Kg^-1^) was decreased compared to WT (**Fig. 7A**). In *Gnal*-D1cKO-/-mice, the progressive decrease in locomotor activity observed after vehicle treatment was prevented, leading to an apparent increase in locomotor activity (**Fig. 7A**). As expected, we observed a strong locomotor hyperactivity in response to 2.5 mg.Kg^-1^ D-amphetamine in WT animals (**Fig. 7B**). With this high dose, large responses were also observed in *Gnal*-D1cKO +/-and *Gnal*-D1cKO-/6 animals. Methylphenidate (10 mg.Kg^-1^) increased locomotor activity similarly in WT and *Gnal*-D1cKO+/-(**Fig. 7C**). The response appeared to be slightly more prolonged in WT than in heterozygotes. In contrast, responses to methylphenidate were completely absent in *Gnal*-D1cKO-/-mice (**Fig. 7C**). These results show that *Gnal* KO in D1-SPNs had contrasted effects on locomotor responses to psychostimulants depending on the drug used. They provide evidence that the D1R/Gα_olf_-dependent activation of AC contributes to the locomotor responses to psychostimulants but that other mechanisms are also involved.

**Fig. 7.**
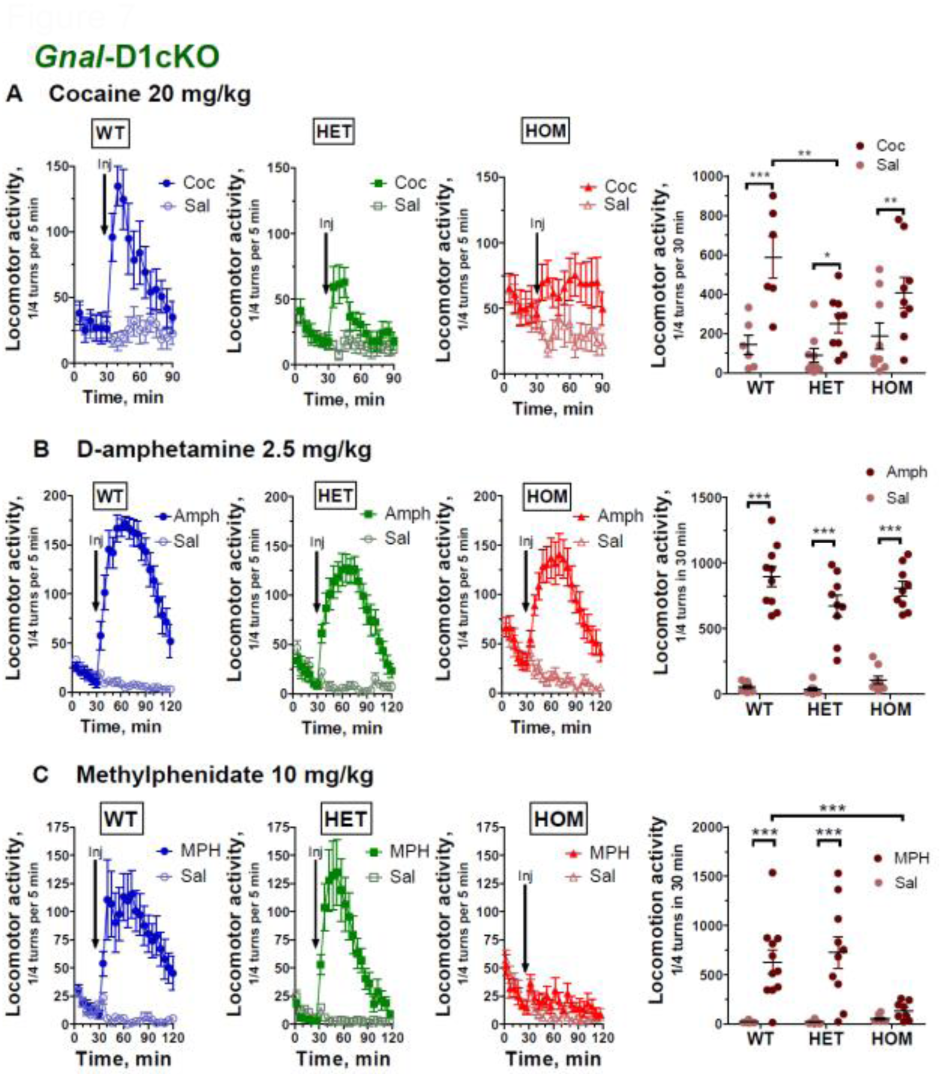
Effects of psychostimulants on locomotion in *Gnal*-D1cKO-/- and *Gnal*-D1cKO+/-mice. **A-C)** time course of locomotor activity (3 left panels) before and after an ip injection of cocaine (Coc, 20 mg.Kg-1) (**A**), D-amphetamine (Amph, 2.5 mg.Kg-1) (**B**), or methylphenidate (MPH, 0.5 mg.Kg-1) (**C**) in wild-type (WT), *Gnal*-D1cKO+/-(HET) and *Gnal*-D1cKO-/-(HOM) mice. In all cases, comparisons were made with time courses after vehicle administration. Each point is the mean ± SEM of the number of ¼ turns in a circular corridor during 5 min. Time-courses of responses to drug and vehicle were compared using repeated measures two-way ANOVA (Treatment and Time factors) followed by Bonferroni’s multiple comparison test (see **Supplementary Statistical tables**). **A-C**, right panels, cumulative responses on 30 min after drug or vehicle administration in WT, HET and HOM mice. Graphs represent scatter dot plots with mean ± SEM. Responses to drug and vehicle in mouse groups with different genotypes were compared using repeated measures two-way ANOVA (Genotype and Treatment factors) followed by Bonferroni’s multiple comparison test (see **Supplementary Statistical tables**). * p<0.05, ** p<0.01, *** p<0.001.

### 3.8 *Gnal* KO in A2A-SPNs alters responses to A2AR antagonists

The A2AR agonist CGS21680 (0.5 mg.Kg^-1^) significantly reduced locomotion in mice with the three different genotypes, including the *Gnal*-A2AcKO-/-mutants which displayed a strong basal hyperlocomotion (**Fig. 8A**). The A2A antagonist KW6002 (3.3 mg.Kg^-1^) increased locomotor activity in WT animals (**Fig. 8B**). This effect was diminished in *Gnal*-A2AcKO heterozygotes and absent in *Gnal*-A2AcKO homozygotes. Caffeine is an antagonist of adenosine receptors with a fairly high affinity, but not selective for A2AR (Fredholm et al., 1999). In WT mice, caffeine (15 mg.Kg^-1^) increased the locomotor activity (**Fig. 8C**). This effect was markedly diminished in *Gnal*-A2AcKO+/-. Interestingly, *Gnal*-A2AcKO-/-mice showed a paradoxical effect following caffeine administration, with a reduction in the basal hyperactivity of these mice. Finally, *Gnal*-A2AcKO-/-mice that were highly hyperactive prior to injection also exhibited a paradoxical hypoactive response following administration of the D1 agonist SKF81297 (5 mg.Kg^-1^) whereas WT and *Gnal*-A2AcKO+/-mice exhibited a hyperactive response (**Fig. 8D**). These results revealed a paradoxical effects of several drugs (caffeine and SKF81297) which increase locomotor activity in WT mice but decreased the spontaneous hyperactivity of *Gnal*-A2AcKO-/-mice.

**Fig. 8.**
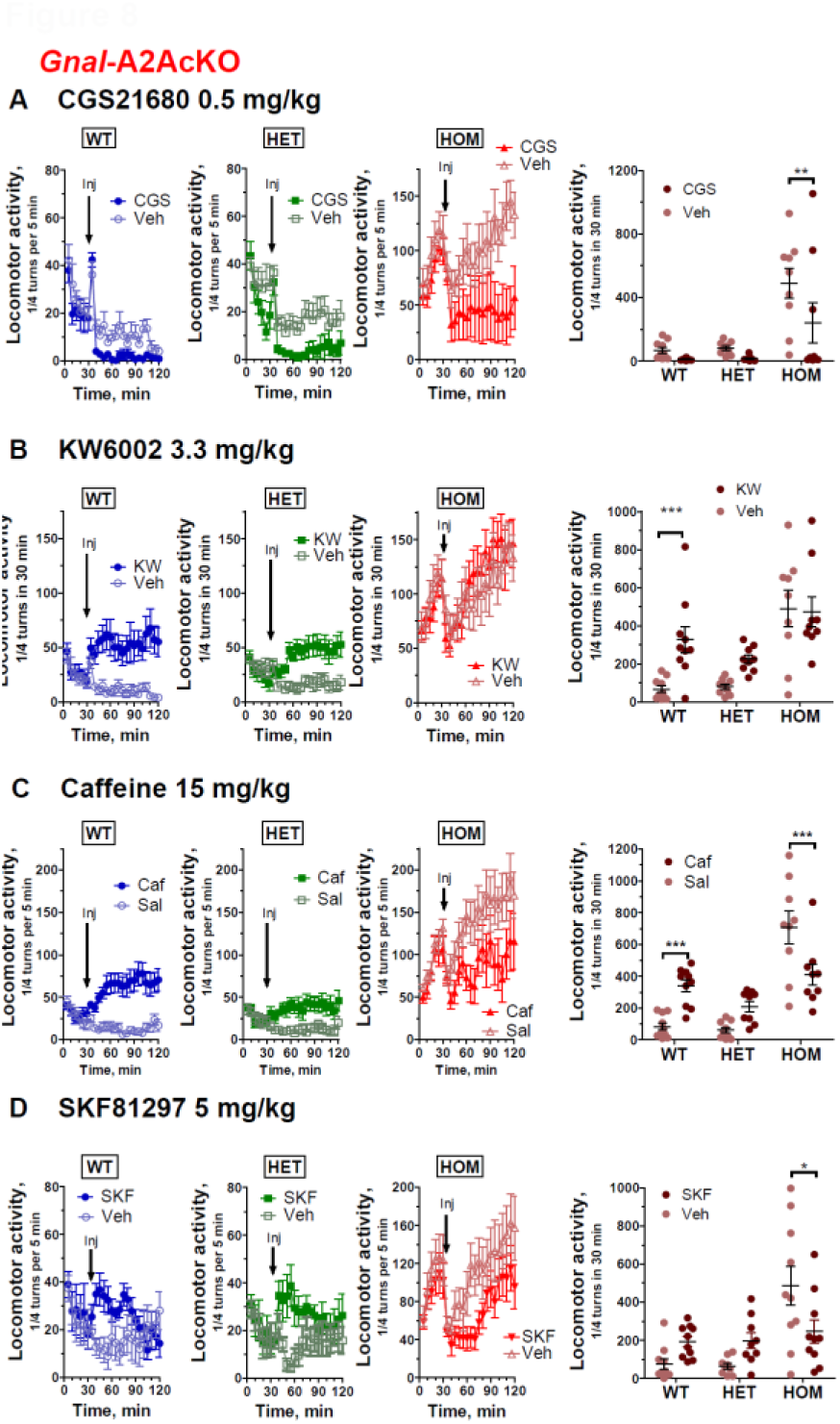
Effects of D1R or A2AR ligands on locomotion in *Gnal*-A2AcKO-/- and *Gnal*-A2AcKO+/- mice. **A-D)** time course of locomotor activity (3 left panels) before and after an ip injection of the A2A agonist CGS21680 (CGS, 0.5 mg.Kg-1) (**A**), A2A antagonist KW6002 (KW or istradefylline, 3.3 mg.Kg-1) (**B**), non-selective adenosine receptor antagonist caffeine (15 mg.Kg-1)(**C**) or D1 agonist SKF81297 (SKF, 5 mg.Kg-1) (**D**) in wild-type (WT), *Gnal*-A2AcKO+/- (HET) and *Gnal*-A2AcKO+/-(HOM) mice. In all cases, comparisons were made with time courses after vehicle administration. Each point is the mean ± SEM of the number of ¼ turns in a circular corridor during 5 min. Time-courses of responses to drug and vehicle were compared using repeated measures two-way ANOVA (Treatment and Time factors) followed by Bonferroni’s multiple comparison test (see **Supplementary Statistical tables**). **A-D**, the right panels, cumulative responses on 30 min after drug or vehicle administration in WT, HET and HOM mice. Graphs represent scatter dot plots with mean ± SEM. Responses to drug and vehicle in mouse groups with different genotypes were compared using repeated measures two-way ANOVA (Genotype and Treatment factors) followed by Bonferroni’s multiple comparison test (see **Supplementary Statistical tables**). * p<0.05, ** p<0.01, *** p<0.001.

### 3.9 *Gnal* KO in A2A-SPNs occludes psychostimulant effects

The hyper-locomotor effects of cocaine (20 mg.Kg^-1^) were similar between WT and *Gnal*-A2AcKO+/-mice (**Fig. 9A**). However, in *Gnal*-A2AcKO-/-mice that were already hyperactive at baseline, no further increase in locomotion was observed following cocaine administration. Similar results were observed in response to D-amphetamine (2.5 mg.Kg^-1^, **Fig. 9B**) or to methylphenidate (10 mg.Kg^-1^, **Fig. 9C**) with comparable locomotor activation in WT and *Gnal*-A2AcKO+/-mice but no additional hyper-locomotor effects in *Gnal*-A2AcKO-/-mice. These results were consistent for the different psychostimulants. Hence, haplo-deficiency of *Gnal* in A2A-SPNs did not modify the response to psychostimulants. In contrast, the locomotor hyperactivity resulting from the absence of *Gnal* expression could not be further increased by psychostimulants, suggesting an occlusion mechanism.

**Fig. 9.**
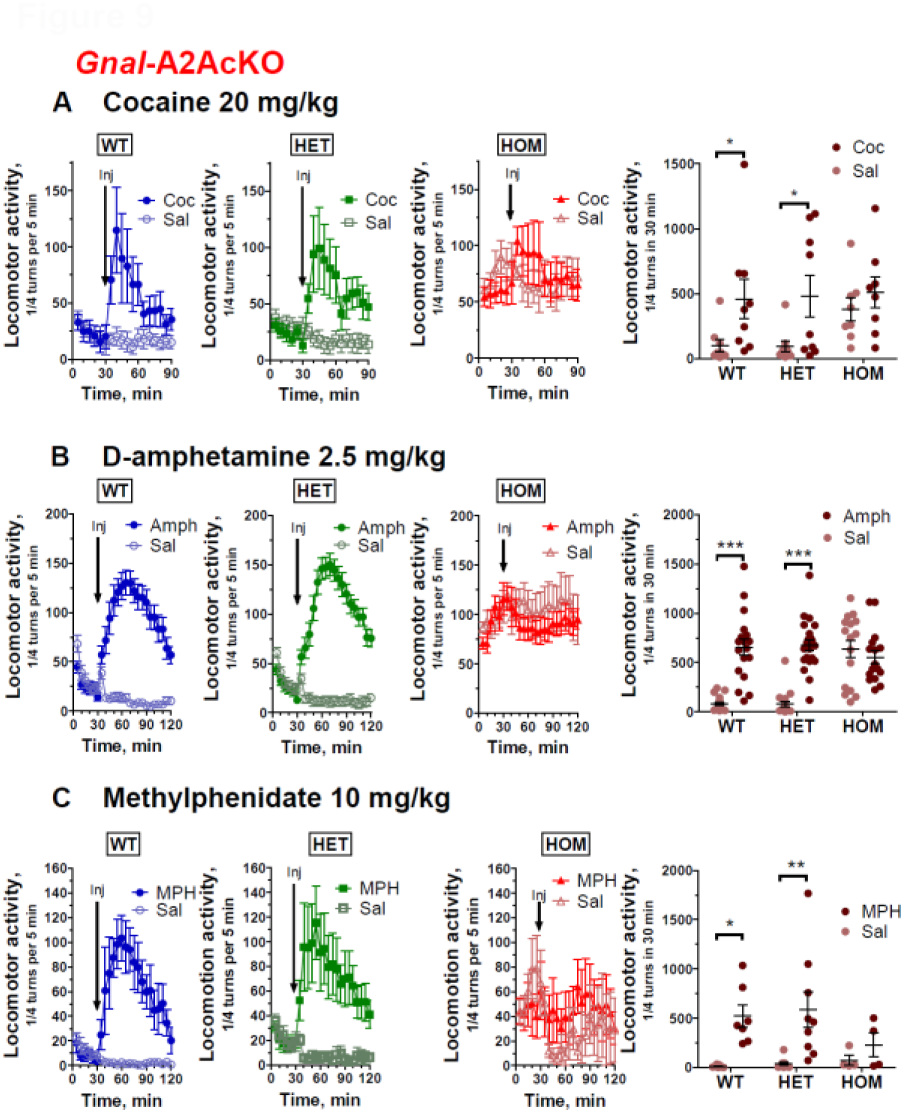
Effects of psychostimulants on locomotion in *Gnal*-A2AcKO-/- and *Gnal*-A2AcKO+/- mice. **A-C)** time course of locomotor activity (3 left panels) before and after an ip injection of cocaine (Coc, 20 mg.Kg-1) (**A**), D-amphetamine (Amph, 2.5 mg.Kg-1) (**B**), or methylphenidate (MPH, 0.5 mg.Kg-1) (**C**) in wild-type (WT), *Gnal*-A2AcKO+/-(HET) and *Gnal*-A2AcKO-/-(HOM) mice. In all cases, comparisons were made with time courses after vehicle administration. Each point is the mean ± SEM of the number of ¼ turns in a circular corridor during 5 min. Time-courses of responses to drug and vehicle were compared using repeated measures two-way ANOVA (Treatment and Time factors) followed by Bonferroni’s multiple comparison test (see **Supplementary Statistical tables**). **A-C**, in the right panels, cumulative responses on 30 min after drug or vehicle administration in WT, HET and HOM mice. Graphs represent scatter dot plots with mean ± SEM. Responses to drug and vehicle in mouse groups with different genotypes were compared using repeated measures two-way ANOVA (Genotype and Treatment factors) followed by Bonferroni’s multiple comparison test (see **Supplementary Statistical tables**). * p<0.05, ** p<0.01, *** p<0.001.

## 4. Discussion

In this study we used conditioned genetic manipulation to dissect the functions of the Gα_olf_ protein, which, together with Gβ_2_ and Gγ_7_, forms the majority of the heterotrimeric G protein that stimulates cAMP production in striatal principal neurons (Schwindinger et al., 2010; Xie et al., 2015). Targeted deletion of *Gnal* (the gene encoding Gα_olf_) abolishes cAMP production stimulated by D1R or A2AR activation in the concerned SPN subpopulation. The absence of Gα_olf_ expression in D1-SPNs or A2A-SPNs results in differential behavioral effects. *Gnal*-D1cKO induces a significant loss of motor performance with mild locomotor hyperactivity, whereas the absence of Gα_olf_ in A2A-SPNs makes mice very hyperactive without significant loss of motor coordination. These effects observed in a rodent isolate the consequences of respective dysfunction in the two populations of SPNs, whose combination might contribute to dystonia in patients. In addition, we report that both *Gnal*-D1cKO and *Gnal*-A2AcKO mice have abnormal responses to drugs interfering with dopamine or adenosine transmission.

### 4.1 Gαolf role in SPNs

The absence of Gα_olf_ in D1-SPNs or A2A-SPNs results in a marked reduction in AC5 levels in the striatum, supporting results obtained in heterozygous constitutive *Gnal* disruption (Xie et al., 2015). These effects mirror those observed after AC5 deletion, leading to a pronounced reduction in Gα_olf_ levels in the striatum (Iwamoto et al., 2004; Xie et al., 2015). *In vivo*, AC5 has been shown to form a stable complex with the regulatory G protein subunits Gα_olf_/β_2_/γ_7_ (Herve, 2011; Schwindinger et al., 2010; Xie et al., 2015) even in the absence of stimulation. The AC5-Gα_olf_ complex has been proposed to form a pre-coupled “signalosome” stabilized by a PLP1 chaperone protein (Xie et al., 2015) in which AC5 activation upon GPCR stimulation results from rearrangement of different subunits rather than their dissociation. The formation of this oligomeric complex protects its various components from proteolysis, an effect that likely allows their precise stoichiometry in the cell. As a result, the absence of Gα_olf_ or AC5 expression increases the degradation of their partner and strongly reduces their cellular concentrations (Herve, 2011). In mice with a *Gnal* cKO, we observed a strong reduction in basal AC activity that may result from the decrease in striatal AC5. Gγ_7_ may play a critical role in the hierarchical assembly of the Gα_olf_/β_2_/γ_7_ complex, because Gγ_7_ is required for the stabilization of Gα_olf_ or Gβ_2_ (Schwindinger et al., 2010) but the presence of Gα_olf_ is not required for Gγ_7_ or Gβ_2_ stabilization. This is consistent with a model in which heterotrimer formation begins with the interaction of Gγ with Gβ to form a stable Gβγ dimer that then associates with an α subunit (Brunori et al., 2021; Pelletier et al., 2024). As a result, the absence of Gα_olf_ expression leaves Gβ_2_/γ_7_ levels unchanged in cells and Gβγ dimer-dependent signaling may remain efficient or be constitutively active due to the loss of control in the heterotrimeric complex. This may contribute to the phenotype of the mutant mice and requires further exploration.

The *Drd1*Cre line has been shown to recombine floxed genes in D1-SPNs (Bateup et al., 2010; Brunori et al., 2021; Lobo et al., 2010). This line recombines in other regions expressing D1R, including the cortex, hippocampus, olfactory bulb, thalamus, but not the cerebellum (Gong et al. 2007; Brunori et al. 2021). There is therefore a possibility that Gα_olf_ disappears in these regions and that this effect contributes to the phenotype of *Gnal*-D1cKO-/- or *Gnal*-D1cKO+/-mice. However, in the olfactory bulb Gα_olf_ levels were not reduced in the mutant mice. In the cortex, despite the significant expression of *Gnal* mRNA (Millett et al., 2024), we were unable to detect Gα_olf_ by western blotting. Moreover, in a recent translatome analysis of D1R-expressing neurons in the cortex, we found that Gα_s_-encoding mRNAs were 5.5 times more abundant than *Gnal* mRNAs (Montalban et al., 2022). This high expression of Gα_s_ would likely compensate for the absence of Gα_olf_. In the cerebellum, D1R expression has only been reported in Bergman glial cells (Cutando et al., 2021), unlike Gα_olf_ which is expressed in Purkinje cells (Belluscio et al., 1998; Millett et al., 2024; Vemula et al., 2013). It is therefore unlikely that Gα_olf_ expression is affected in the cerebellum. In the Adora2aCre line, it has been shown that genetic recombination is highly selective for SPNs expressing enkephalin (Durieux et al., 2009). The piriform cortex is the only region where recombinant neurons have been observed (Durieux et al., 2009).

In the striatum, selective *Gnal*-D1cKO results in complete loss of response to the D1 agonist SKF81297, demonstrating a necessary role for Gα_olf_ in coupling D1R to AC. Gα_s_ is also expressed in the striatum, mainly in cholinergic and GABAergic interneurons and in afferent fibers arriving to the striatum but very little in SPNs (Herve et al., 2001; Millett et al., 2024; Montalban et al., 2022). The absence of response to the A2A agonist CGS21680 in *Gnal*-A2AcKO-/-mice demonstrates that Gα_olf_ is also essential for coupling A2AR to AC. In the striatum of *Gnal*-D1cKO-/-mice, the response of AC to an A2AR agonist was not significantly altered and in *Gnal*-A2AcKO-/-mice, AC response to a D1R agonist remained unchanged. These results are consistent with the overall dichotomy of SPNs in the dorsal striatum (Gerfen and Surmeier, 2011), in spite of the existence of a minor population expressing both D1R and D2R in rodents (Bertran-Gonzalez et al., 2008; Bonnavion et al., 2024).

### 4.2 *Gnal* deletion in D1-SPN impairs motor abilities

The absence of Gα_olf_ expression in D1-SPNs results in poor performances in the Rotarod and pole tests. This phenotype cannot be attributed to a generalized motor deficit since the mutant mice exhibit high locomotor activity especially during the night and do not show any weakness. Mutant mice did not show any sign of abnormal risk assessment that could bias the results in the Rotarod or pole tests. In the Rotarod, *Gnal*-D1cKO-/-mice are capable of motor learning with the repetition of training sessions, but learning is delayed compared to WT mice. Mice with constitutive deletion of D1Rs show similar deficits in the Rotarod (Karasinska et al., 2005; Kobayashi et al., 2004; Short et al., 2006; Wall et al., 2011) in the first days of testing, but are also able to increase their performance after multiple learning sessions (Nakamura et al., 2014). Mice lacking AC5 expression also show reduced motor performance in the Rotarod and pole test (Iwamoto et al., 2003) while some motor learning can still be observed. Together these data support an important role of the D1R/Gα_olf_/AC5 complex in D1-SPNs in motor performances. This motor deficit may be partially compensated by other neural circuits and/or intracellular signaling pathways, allowing learning after prolonged training. Poor performance at in the Rotarod test with apparently aberrant motor learning have been observed in other animal models of dystonia (reviewed in (Richter and Richter, 2014)). This phenotype may represent a consistent species-specific modeling of human dystonic symptoms in mice. In contrast, *Gnal*-A2AcKO-/-mice do not show any impairment in motor performance or learning in the Rotarod test. This is consistent with data showing that ablation of A2A-SPNs has less effect in the Rotarod than ablation of D1-SPNs (Durieux et al., 2012).

### 4.4 *Gnal* knockout in D1-SPN impairs responses to drugs

*Gnal*-D1cKO has contrasting effects on locomotor responses to various D1 agonists. Responses to SKF81297 or dihydrexidine are completely or largely abolished whereas SKF83822-induced hyperactivity is preserved. Contrasting alterations are also observed in responses to cocaine, D-amphetamine, and methylphenidate. *Gnal*-D1cKO-/-mice no longer respond to methylphenidate while they exhibit decreased responses to cocaine and virtually no alteration in D-amphetamine effects. It is therefore clear that part of the locomotor response to these drugs depends on the presence of Gα_olf_ in D1-SPNs, but that another part does not, the proportion between the two varying from one drug to another. Responses to SKF81297 or cocaine are largely mediated by dopamine D1Rs, since they are completely abolished by disruption of *Drd1* gene (Xu et al., 1994a; Xu et al., 1994b). In contrast, the absence of D1Rs only attenuates responses to D-amphetamine (Crawford et al., 1997; Xu et al., 2000). Deletion of Gγ7 in D1-SPNs, which strongly reduces Gα_olf_ expression in the striatum, does not significantly alter locomotor responses to the D1R agonists SKF83822 or SKF83959 as well as to D-amphetamine (Brunori et al., 2021). As discussed above, it is unlikely that Gα_s_ is responsible for the partly preserved responses to D1 agonists in *Gnal*-D1cKO-/-mice. D1R activates several cAMP-independent signaling pathways, involving β-arrestins (Conroy et al., 2015; Kaya et al., 2020; Urs et al., 2011), G_q_/Phospholipase C (Jin et al., 2003; Lee et al., 2014; Medvedev et al., 2013; Undie and Friedman, 1990), tyrosine kinase Src leading to NMDA receptor phosphorylation (Kaya et al., 2020; Pascoli et al., 2011) and D1R heteromerization with NMDA receptor subunits (Andrianarivelo et al., 2021; Cahill et al., 2014; Fiorentini et al., 2003; Lee et al., 2002). All of these alternative signaling pathways could mediate responses to D1R-stimulating drugs in the absence of Gα_olf_. In addition, different D1R agonists exhibit biases in their abilities to activate different signaling pathways downstream of D1Rs (Conroy et al., 2015; Jin et al., 2003; Jones-Tabah et al., 2021; Martini et al., 2019; Yano et al., 2018) which could be a factor explaining different consequences of Gα_olf_ absence in D1-SPNs.

### 4.5 *Gnal* KO in A2A-SPN induces a high locomotor hyperactivity

*Gnal* disruption in A2A-SPNs leads to a strong spontaneous locomotor hyperactivity, particularly during the nocturnal phases corresponding to the waking period in mice, during which the locomotor activity of *Gnal*-A2AcKO-/-mice was ∼15-fold higher than in WT animals. Importantly, this hyperactivity is prolonged and even tends to increase in a familiar environment, unlike what is observed in WT mice. This hyperactivity is similar to that observed when A2A-SPNs, particularly those in the dorsomedial region of striatum, are selectively destroyed by diphtheria toxin (Durieux et al. 2009; Durieux et al., 2012). However, *Gnal*-A2AcKO-/-mice do not show any evidence of A2A-SPN degeneration since the levels of DARPP-32, a specific marker of SPNs, are normal in these mice. Spontaneous locomotor hyperactivity is also observed when Gγ7 or DARPP-32 is lacking in A2A-SPNs (Bateup et al., 2010; Brunori et al., 2021). This locomotor hyperactivity could be the consequence of disrupted A2AR activity. However, complete A2AR deletion or selective A2AR deletion in the striatum does not significantly alter spontaneous locomotor activity (Ledent et al., 1997; Shen et al., 2008). Although A2AR is probably the most abundantly expressed receptor stimulating AC in A2A-SPNs other receptors may compensate for its absence. It is also possible that hyperlocomotion is related to decrease of basal cAMP production, rather than impairment of cAMP-dependent signaling of A2AR. In these mice, the selective A2AR antagonist KW6002 (istradefylline) loses its hyperlocomotor effect while caffeine decreases locomotion, results reminiscent of those observed in mutant mice lacking A2AR (Ledent et al., 1997) or Gγ7 in A2A-SPNs (Brunori et al., 2021). Interestingly, the effects of KW6002 and caffeine are reduced in *Gnal*-A2AcKO+/-mice compared to WT mice, confirming similar results in mice with constitutive *Gnal* hemizygoty (Herve et al., 2001). This clearly shows that the psychostimulant effects of caffeine and KW6002 are mediated by the decrease in A2AR-dependent cAMP production. In the absence of Gα_olf_ in A2A-SPNs, all psychostimulant drugs tested lose their effects on locomotor activity. The D1R agonist, SKF81297 even acquires a hypo-locomotor effect. However, other effects of SKF81297 such as the induction of epileptic seizures (Gangarossa et al., 2011) could disrupt locomotor activity in mutant mice. Given the high baseline hyperactivity of these mice, a ceiling effect cannot also be excluded, blocking any further increase in locomotor activity.

### 4.6 *Gnal* KO in D1-SPNs increases mortality around weaning

We observed an abnormally high mortality rate of *Gnal*-D1cKO-/-mice around weaning. Approximately a third of homozygotes died during this period. Those mice ceased gaining weight around day 15, a period when mothers gradually stop nursing their pups. The other remaining mutant mice were apparently not affected during this period with a normal development and a body weight similar to that of WT mice as adults. The most likely hypothesis is that the gradual transition from milk feeding by the mother to a more solid and diverse diet (weaning) is disrupted in some mutant mice. In mice with a constitutive deletion of *Gnal*, a very high newborn lethality was also observed at an earlier time, from the first days after birth. It is generally accepted that this mortality is due to the early anosmia of these mice, caused by a loss of olfactory receptor signaling (Belluscio et al., 1998). The pups would no longer detect the olfactory signals necessary for suckling. However, Gα_olf_ deficits in striatal neurons could also contribute to the lethality in pups since very high infant mortality has been reported in a model of conditioned deletion in all striatal neurons from the embryonic stages (Chambers et al., 2024). Mutant mice with severe dopamine deficiency that were indistinguishable from their WT littermates by postnatal week 1, became hypoactive and stopped feeding by postnatal days 10-15 (Zhou and Palmiter, 1995). A similar phenotype and lethality were observed in mice lacking ERK1/2 in the D1-SPNs (Hutton et al., 2017). In our *Gnal*-D1cKO, mortality was less massive affecting only 1/3 of mutant mice but the developmental halt appeared at the same period as that observed in dopamine or ERK1/2-deficient mice. This P10-P15 period corresponds to a time when both Gα_olf_ and AC5 expression increase sharply in the developing striatum (Iwamoto et al., 2004; Rius et al., 1994). Gα_olf_ expression is very low at birth and reaches its maximum at P21 corresponding to weaning. At birth, Gα_s_ expression exceeds that of Gα_olf_ but the ratio reverses during the second postnatal week. It is therefore likely that functional D1-SPN disruption only begins to be significant from this second week, possibly impairing motivational and learning processes important for feeding. Our results suggest that D1-SPNs play an important role in these behavioral adaptations at weaning. The reason why this lethal phenotype and the prior loss of weight appeared in only a third of the mice remains to be determined but we could speculate that specific compensatory mechanisms could be insufficient at this critical period of life in a specific part of the pups. It is notable that *Gnal*-A2AcKO-/-mice have normal viability, revealing a non-critical role of A2A-SPNs around weaning.

### 4.6 Limitations of the study

This study is the first general exploration of *Gnal* cKO examining the effect of the selective deletion of *Gnal* in each of the two specific subpopulations of SPNs. We did not observe spontaneous dystonic movement but further exploration with recording of muscles and the use of drugs potentially triggering dystonia, such as muscarinic agonists (Pelosi et al., 2017) will be important as well as characterization of a potential dysfunctions of the cerebello-thalamic pathways (Aissa et al., 2022). Concerning the exploration of responses to pharmacological agents, the use of dose-response curve and blockade of responses with specific antagonists would be informative. Because of their high hyperactivity, *Gnal*-A2AcKO mice could be a mouse model of ADHD. However, its validation would require determining whether these mice exhibit attention disturbances and whether chronic treatment with methylphenidate and amphetamine reduces hyperactivity and possible attention deficits.

### 4.7 Conclusions

Our study shows that Gα_olf_-mediated striatal signaling in selective populations of SPNs differentially controls motor behavior and drug responses in mice. Our results confirm that Gα_olf_ is critical for the activation of cAMP production induced by stimulation of D1R in D1-SPNs and A2AR in A2A-SPNs. Gα_olf_/β2/γ7 heterotrimers stabilize AC5 signalosome (Herve, 2011; Xie et al., 2015). We propose that the observed phenotypes arise from the destabilization of this signalosome, leading to decreased basal cAMP production and a loss of receptor-mediated stimulation, particularly affecting D1R and A2AR. Gα_olf_ deficiency in D1-SPNs induces significant disturbances in motor control that could promote *GNAL*-related dystonic disorders in humans. In contrast, when Gα_olf_ is lacking in A2A-SPNs, mice become highly hyperactive without overt motor coordination impairment. This hyperactivity may result from a disinhibition of movement which might also contribute to the development of dystonic symptoms in patients with DYT-*GNAL*. The hypothesis of a joint involvement of D1-SPNs and A2A-SPNs is consistent with findings of a recent study. Selective suppression of *Gnal* in the striatum, affecting both D1-SPNs and A2A-SPNs, but also the other types of striatal neurons), induces abnormal movements in mice, reminiscent of human dystonic symptoms (Chambers et al., 2024). The phenotype is more pronounced when *Gnal* KO occurs in adulthood using a viral approach. One possible explanation is that these mice cannot engage in compensatory processes during the development of the motor system to mitigate the consequence of the striatal dysfunction associated with GNAL suppression. However, in this study, *Gnal* disruption affects all types of striatal neurons, notably cholinergic neurons and parvalbumin-positive GABAergic neurons that express Gα_olf_ in addition to Gα_s_ (Herve et al., 2001; Millett et al., 2024). The absence of Gα_olf_ in these cholinergic and GABAergic neurons may also contribute to the development of the observed dystonic-like phenotype.

Interestingly, locomotor responses to D1 agonists and psychostimulants are differentially affected by the absence of Gα_olf_ in D1-SPNs, showing that these responses are only partially dependent on Gα_olf_-dependent D1R signaling. Our results suggest the possibility that alternative cAMP-independent D1R-effectors contribute to the effects of psychostimulant drugs. Thus, we identify Gα_olf_ as an important, but not exclusive, mediator of D1R and A2AR signaling, which would make it an interesting target for the development of modulatory drugs as new therapeutic options for dystonia but also other neurological or psychiatric diseases, such as Parkinson’s disease and drug addiction.

## Supporting information

Supplementary Statistical tables

## Abbreviations

A2AR: A2A receptor
AC: adenylyl-cyclase;
AC5: adenylyl-cyclase 5;
D1R: D1 receptor;
D2R: D2 receptor;
DARPP-32: dopamine and cAMP-regulated phospho-protein 32 kDa;
DYT-GNAL: dystonia linked to *GNAL* gene;
*Gnal*-A2AcKO: *Gnal* conditional knockout in A2A receptor-expressing neurons;
*Gnal*-D1cKO: *Gnal* conditional knockout in D1 receptor-expressing neurons;
GPCR: G protein-coupled receptor;
HET: heterozygous,
HOM: homozygous;
ip: intraperitoneal;
KO: knockout;
SPN: striatal projection neuron;
A2A-SPN: A2A receptor-expressing striatal projection neuron;
D1-SPN: D1 receptor-expressing striatal projection neuron;
dSPN: striatal projection neuron of direct pathway;
iSPN: striatal projection neuron of indirect pathway;
WT: wild-type

## CRediT authorship contribution statement

Sophie Longueville: Formal analysis, Investigation, Methodology, Writing. Ruiyi Yuan: Investigation, Methodology. Claire Naon: Investigation, Methodology. Emmanuel Valjent: Data curation, Formal analysis, Investigation, Methodology, Writing. Assunta Pelosi: Investigation, Methodology, Writing. Emmanuel Flamand-Roze: Resources, Funding acquisition, Supervision, Writing. Louise-Laure Mariani: Resources and Funding acquisition, Investigation, Methodology, Supervision, Writing. Jean-Antoine Girault: Conceptualization, Formal analysis, Funding acquisition, Project co-administration, Resources, Supervision, Writing. Denis Hervé: Formal analysis, Investigation, Methodology, Conceptualization, Data curation, Formal analysis, Funding acquisition, Project administration, Resources, Supervision, Validation, Writing.

## Data availability

Data will be made available on request.

## Acknowledgments

We thank the *Mouse breeding and phenotyping facility* for animal care of the Institut du Fer à Moulin (IFM). Part of the animal work was conducted at the PBI Pheno-ICMice Core Facility. The Core is supported by 2 “Investissements d’avenir” (ANR-10-IAIHU-06 and ANR-11-INBS-0011-NeurATRIS) and the “Fondation pour la Recherche Médicale”. We gratefully acknowledge Nadege Sarrazin for assistance with animal behavior facility use. This work benefited from equipment and services from the iGenSeq core facility (Genotyping and Sequencing) at PBI (Paris Brain Institute).

