## Supplementary figures_S1-S6 for "Characterization of mice with cell type-specific *Gnal* loss of function provides insights on *GNAL*-linked dystonia"

Fig. S1

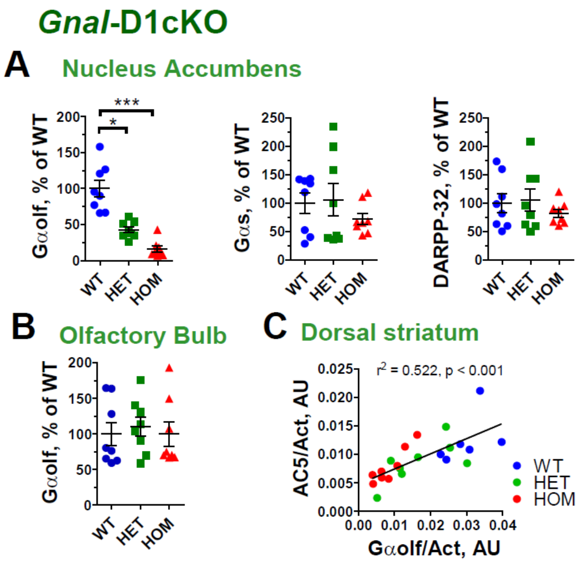


#### Fig. S1 Molecular characterization of *Gnal*-D1cKO-/- and *Gnal*-D1cKO+/- mice.

**A)** Quantification of $G\alpha_{olf}$, $G\alpha_s$ and DARPP-32 levels in *nucleus accumbens* homogenates of *Gnal*-D1cKO-/- (HOM), *Gnal*-D1cKO+/- (HET) and wild-type (WT) littermates. Partial or complete loss of *Gnal* expression in D1-SPNs reduced significantly the levels of $G\alpha_{olf}$, but not those of Gαs or DARPP-32. **B)** Quantification of $G\alpha_{olf}$ levels in olfactory bulb homogenates. These levels were not altered in *Gnal*-D1cKO HOM and HET mice compared to WT mice. Data corresponded to the ratio between the immunofluorescence of the indicated protein and actin, expressed as a % of the mean value in WT samples. Protein levels were compared using Kruskal-Wallis test followed Dunn's multiple comparison test (see the results in **Supplementary Statistical tables**). * $p<0.05$, ***, $p < 0.001$. **C)** Linear regression between the $G\alpha_{olf}$ and AC5 levels in dorsal striatum samples of mutant and WT mice ($r^2 = 0.522$, $p<0.001$).

Fig. S2

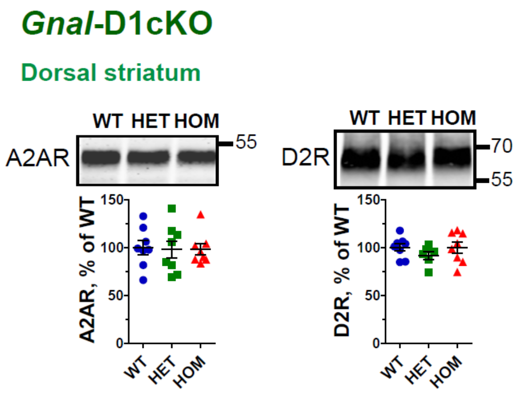


#### Fig. S2 A2A and D2 receptor levels in *Gnal*-D1cKO-/- and *Gnal*-D1cKO+/- mice

Representative immunoblots and quantification of A2A adenosine receptor (A2AR) and D2 dopamine receptor (D2R) in dorsal striatum homogenates of *Gnal*-D1cKO-/- (HOM), *Gnal*-D1cKO+/- (HET) and wild-type (WT) littermates. Data corresponded to immunofluorescence of the indicated protein, expressed as a % of the mean value in WT samples. Protein levels were compared Kruskal-Wallis test followed Dunn's multiple comparison test. No significant changes were observed in mutant mice compared to WT mice (see the results in **Supplementary Statistical tables**).

Fig. S3

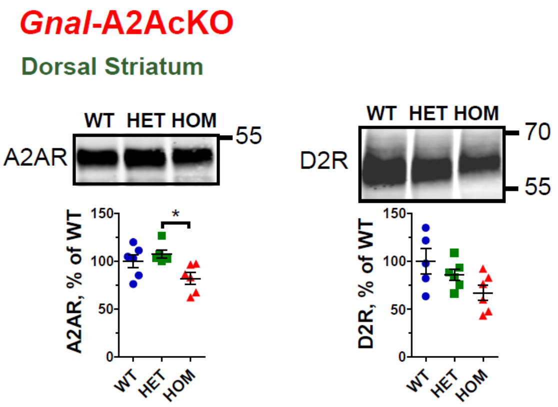


#### Fig. S3 A2A and D2 receptor levels in *Gnal*-A2AcKO-/- and *Gnal*-A2AcKO+/- mice

Representative immunoblots and quantification of A2A adenosine receptor (A2AR) and D2 dopamine receptor (D2R) in dorsal striatum homogenates of *Gnal*-A2AcKO-/- (HOM), *Gnal*-A2AcKO+/- (HET) and wild-type (WT) littermates. Data corresponded to immunofluorescence of the indicated protein, expressed as a % of the mean value in WT samples. Protein levels were compared Kruskal-Wallis test followed Dunn's multiple comparison test. * $p < 0.05$, a significant decrease of A2AR levels was observed in HOM mutant mice compared to WT mice (see the results in **Supplementary Statistical tables**).

Fig. S4

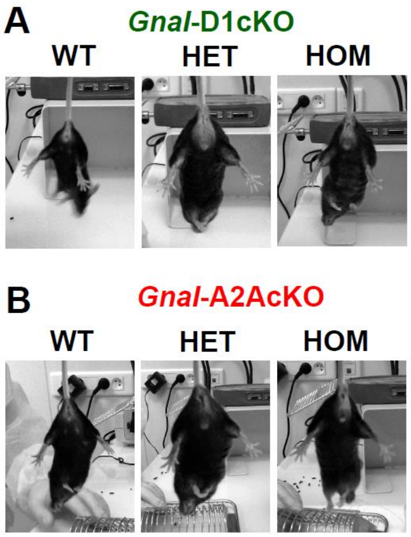


#### Fig. S4 Lack of hindlimb clasping in *Gnal*-D1cKO and *Gnal*-A2AcKO mice

Representative images from videos show the posture of mice when suspended by the tail. Comparison of wild-type (WT), homozygous (HOM) and heterozygous (HET) mutant mice in the *Gnal*-D1cKO (**A**) and *Gnal*-A2AcKO (**B**) lines. Mutant mice in both lines show no hindlimb clasping nor, in general, any overt movement abnormalities.

Fig. S5

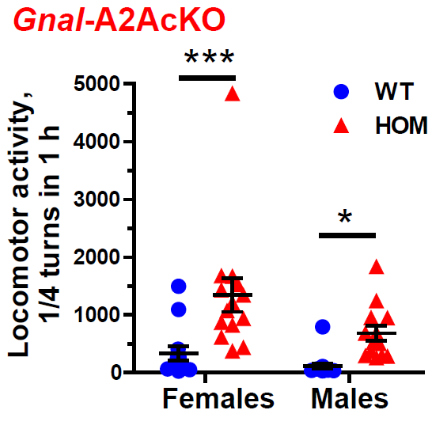


#### Fig. S5 Increase locomotion in female and male *Gnal*-A2AcKO-/- mice.

Locomotor activity during 1 h after saline injection in wild-type (WT) and *Gnal*-A2AcKO-/- (HOM) mice. Comparison of females and males. The animals were habituated to the experimental conditions for 2 consecutive days. On day 3, the animals were placed in the circular maze and, 30 min later, received a saline injection. Recording began after the injection and lasted for 1 h. Females were more active than male. In both sexes, locomotor activity in *Gnal*-A2AcKO-/- was increased compared to WT mice. Graphs represent scatter dot plots with mean ± SEM. Effects of Sex and Genotypes were analyzed using two-way ANOVA followed by Bonferroni's multiple comparison test (see **Supplementary Statistical tables**). * $p<0.05$, *** $p<0.001$.

### Fig. S6

#### *Gnal*-D1cKO

**SKF83822 0.5 mg/kg**

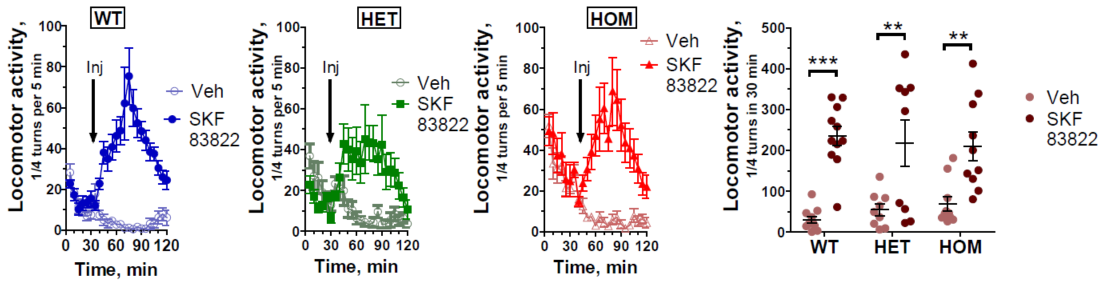


Fig. S6 Effects of the D1R agonist SKF83822 on locomotion in *Gnal*-D1cKO-/- and *Gnal*-D1cKO+/- mice.

In the 3 left panels, time course of locomotor activity, before and after an ip injection of the D1R agonist SKF83822 (0.5 mg.Kg$^{-1}$) in wild-type (WT), *Gnal*-D1cKO+/- (HET) and *Gnal*-D1cKO-/- (HOM) mice. Comparisons were made with locomotion time course after vehicle administration. Each point is the mean ± SEM of the number of ¼ turns in a circular corridor during 5 min. Time-courses of responses to SKF83822 and vehicle were compared using repeated measures two-way ANOVA (Treatment and Time factors) followed by Bonferroni's multiple comparison test (see **Supplementary Statistical tables**). Right panel, cumulative responses on 30 min after SKF83822 or vehicle administration in WT, HET and HOM mice. Graphs represent scatter dot plots with mean ± SEM. Responses to drug and vehicle in mouse groups with different genotypes were compared using repeated measures two-way ANOVA (Genotype and Treatment factors) followed by Bonferroni's multiple comparison test (see **Supplementary Statistical tables**). ** $p<0.01$, *** $p<0.001$.
